# Diminished Immune Cell Adhesion in Hypoimmune ICAM-1 Knockout Pluripotent Stem Cells

**DOI:** 10.1101/2024.06.07.597791

**Authors:** Sayandeep Saha, W. John Haynes, Natalia M. Del Rio, Elizabeth E. Young, Jue Zhang, Jiwon Seo, Liupei Huang, Alexis M. Holm, Wesley Blashka, Lydia Murphy, Merrick J. Scholz, Abigale Henrichs, Jayalaxmi Suresh Babu, John Steill, Ron Stewart, Timothy J. Kamp, Matthew E. Brown

## Abstract

Hypoimmune gene edited human pluripotent stem cells (hPSCs) are a promising platform for developing reparative cellular therapies that evade immune rejection. Existing first-generation hypoimmune strategies have used CRISPR/Cas9 editing to modulate genes associated with adaptive (e.g., T cell) immune responses, but have largely not addressed the innate immune cells (e.g., monocytes, neutrophils) that mediate inflammation and rejection processes occurring early after graft transplantation. We identified the adhesion molecule ICAM-1 as a novel hypoimmune target that plays multiple critical roles in both adaptive and innate immune responses post-transplantation. In a series of studies, we found that ICAM-1 blocking or knock-out (KO) in hPSC-derived cardiovascular therapies imparted significantly diminished binding of multiple immune cell types. ICAM-1 KO resulted in diminished T cell proliferation responses *in vitro* and in longer *in vivo* retention/protection of KO grafts following immune cell encounter in NeoThy humanized mice. The ICAM-1 KO edit was also introduced into existing first-generation hypoimmune hPSCs and prevented immune cell binding, thereby enhancing the overall hypoimmune capacity of the cells. This novel hypoimmune editing strategy has the potential to improve the long-term efficacy and safety profiles of regenerative therapies for cardiovascular pathologies and a number of other diseases.

**Highlights:**

- Antibody blocking of ICAM-1 on human pluripotent stem cell-derived cells inhibits immune cell adhesion
- CRISPR/Cas9 knock-out of ICAM-1 ablates surface and secreted ICAM-1 protein and inhibits immune adhesion
- ICAM-1 knock-out results in decreased T cell proliferative responses to human pluripotent stem cell-derived grafts *in vitro*, and resistance to immune-mediated graft loss *in vivo*
- Addition of ICAM-1 knock-out to first generation MHC knock-out human pluripotent stem cells confers protection against immune adhesion

Graphical Abstract
ICAM-1 Knock-out in Transendothelial Migration and at the Immune Synapse. Abbreviations: PSC-EC – pluripotent stem cell-derived endothelial cells; KO – knock-out; dSMAC – distal supramolecular activation complex; pSMAC – peripheral supramolecular activation complex; cSMAC – central supramolecular activation complex.

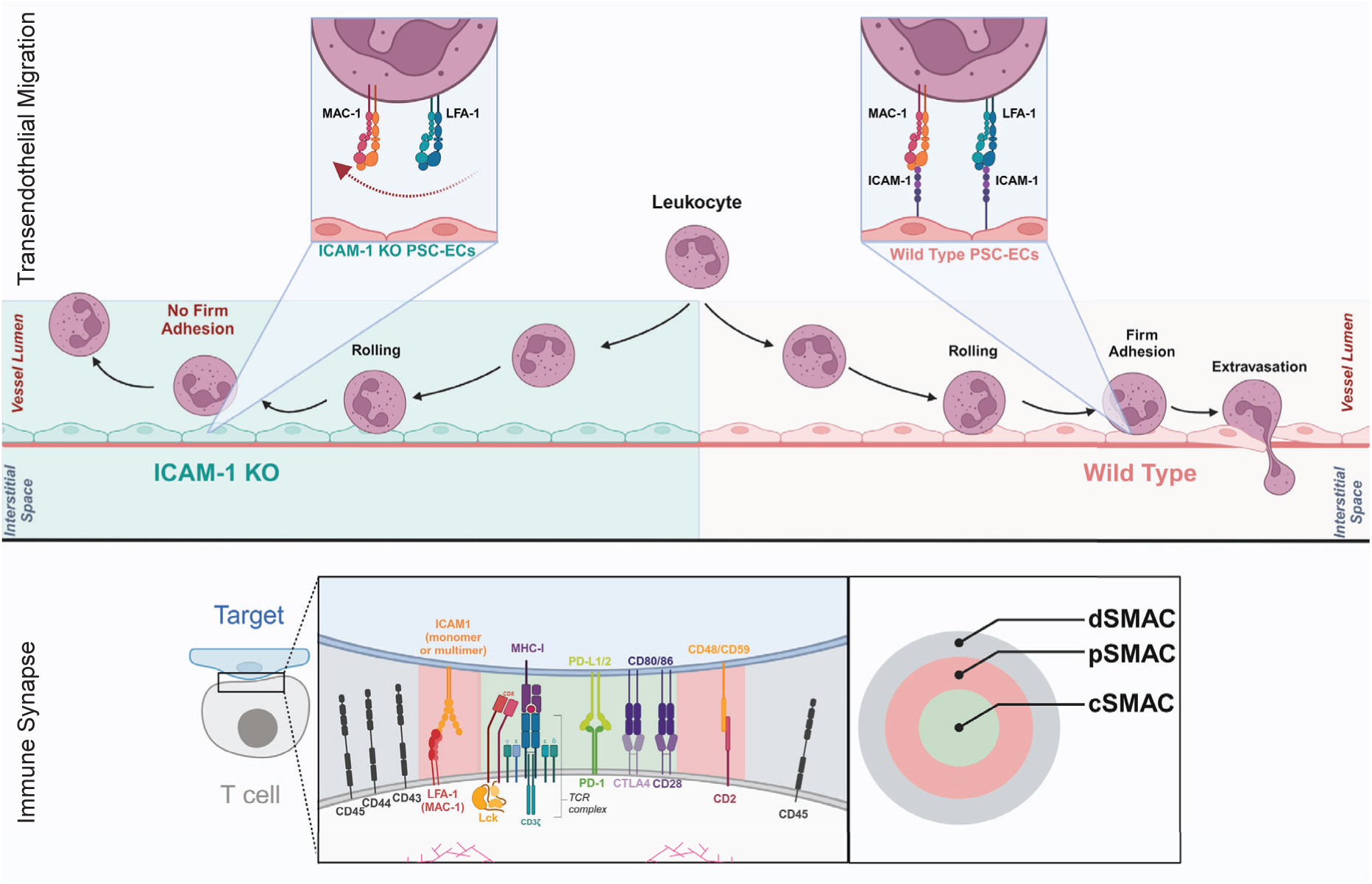

## Introduction

Human pluripotent stem cell (hPSC)^1,2^ technology is a promising cellular therapy platform for treating multiple diseases, including acute cardiovascular pathologies (e.g., myocardial infarction),^3^ Parkinson’s disease,^4^ and autoimmune disorders (e.g., type 1 diabetes).^5^ Two key attributes of hPSCs are: 1) scalability, which allows cells from a single donor to treat potentially thousands of patients; and 2) amenability to gene editing, including “hypoimmune” strategies to modulate genes involved in alloimmune rejection with the goal of achieving immune tolerance and long-term retention of hPSC-based grafts.^6^ The first generation of hypoimmune hPSC gene edits largely focuses on targeting important mechanisms of adaptive immunity-driven allograft rejection, namely T cell-mediated rejection and antibody-mediated rejection. Hypoimmune editing strategies include genetic knock-out (KO) of major histocompatibility complex (MHC) class I and II surface expression,^7,8^ individually and with additional knock-in edits to mitigate natural killer cell-mediated cytotoxicity (an adverse effect resulting from MHC KO).^9,10^ While these adaptive immunity-centric methods are promising, innate immune cells (e.g., neutrophils, monocytes) are intimately involved in the very earliest stages of allorejection of solid organs and cellular therapies^11–15^ and thus remain a critical concern that has not been fully addressed by existing hypoimmune strategies.^16^

We sought to identify novel hypoimmune editing targets by first evaluating genes involved in adaptive as well as innate immune allorejection pathways. Adaptive and innate immune cells express the integrins LFA-1 (CD11a/CD18) and/or MAC-1 (CD11b/CD18) (Fig. 1a, Supplementary Fig. 1a-c), both of which bind to Intercellular Adhesion Molecule 1 (ICAM-1), a glycoprotein expressed on the cell surface and in secreted forms by endothelial cells (ECs), cardiomyocytes (CMs), epithelial cells, fibroblasts, and various immune cells. Immune cell LFA-1 and/or MAC-1-mediated binding to ICAM-1 on ECs is a critical step in the very earliest inflammatory stages of transplant allorejection, as it stabilizes the immune synapse that precedes effector-mediated killing and plays a key role in the firm adhesion process required for extravasation of immune cells into the parenchyma of transplanted grafts.^17^ These binding interactions also directly impact T cell activation^18^ and effector function,^19^ making ICAM-1 an attractive editing target with potentially pleiotropic hypoimmune effects. Previous studies in mice have demonstrated that KO of ICAM-1 isoforms may have a beneficial impact on oxidative stress responses during inflammation^20^ and that absence of surface ICAM-1 diminishes immune cell binding to ECs^21^ and other parenchymal cell types, resulting in prolongation of allograft survival in heart transplant studies.^22^ In humans, clinical studies have assessed targeting ICAM-1 via blocking antibodies^23–25^ and antisense oligonucleotides^26^ in transplantation and other settings with conflicting results, potentially due to the systemic effects of these approaches and/or to the recombinant antibody design strategies used at the time of these early trials, which started over a decade ago. However, clinical investigation with new ICAM-1 blocking antibodies is still an active area of research in the human solid organ^27^ and xenotransplantation^28^ fields.

**Figure 1.**
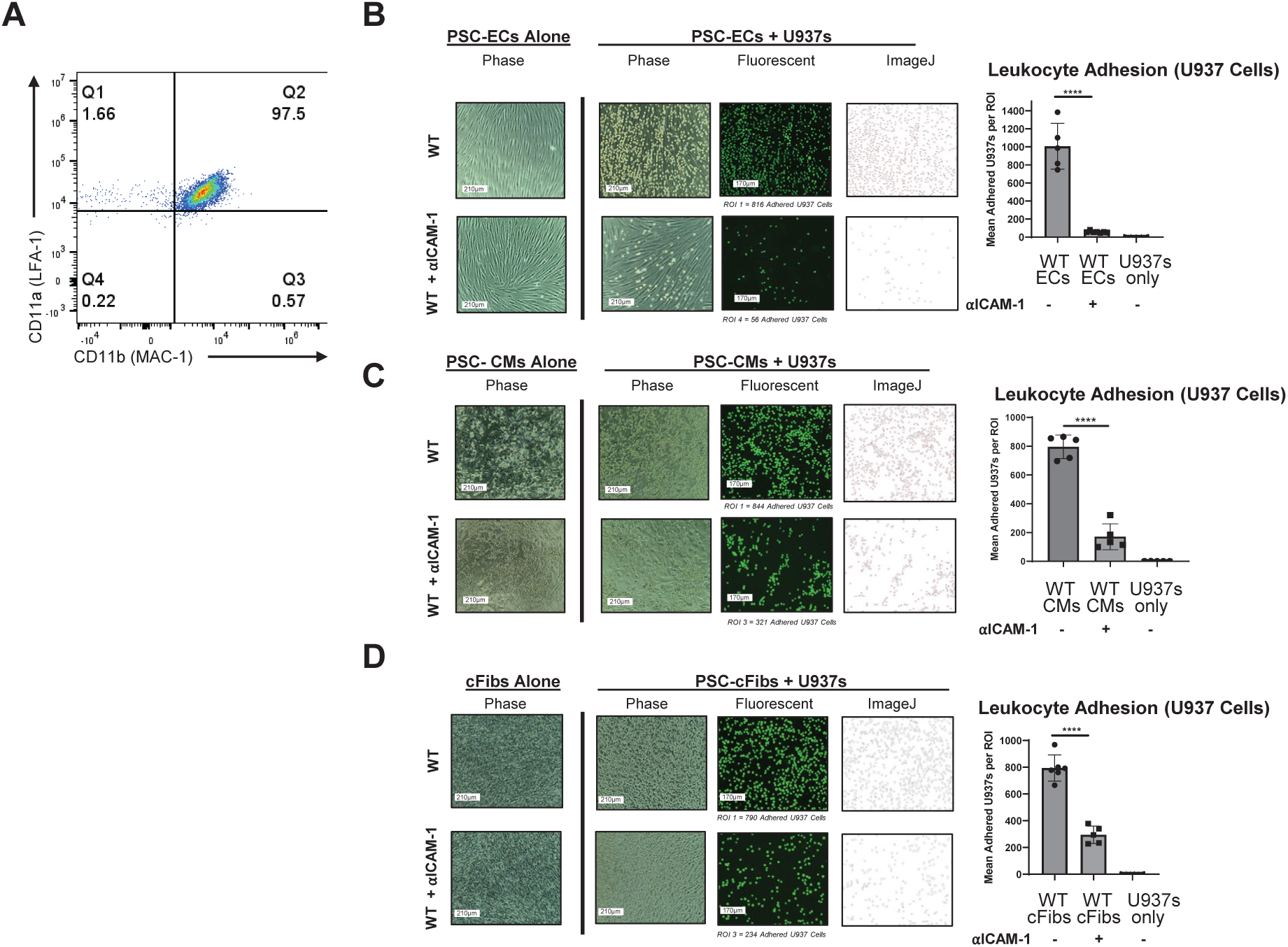
Adhesion Assay with Antibody Blocking of ICAM-1 in Pluripotent Stem Cell (PSC)-Derived Cardiovascular Cells. (A) The U937 monocytic cell line was stained with LFA-1 (CD11a/CD18) and MAC-1 (CD11b/CD18) alpha chain antibodies. (B) PSCs (H9 embryonic stem cell line) were differentiated into high-purity endothelial cells (ECs) (>85% CD31+CD144+), stimulated with TNFα (10ng/ml) and IFNγ (50ng/ml) for 48 hours, followed by incubation with anti-human ICAM-1 blocking antibody (αICAM-1) (0.5mg/ul) for 1 hour at 37°C, and washed. U937 cells (labeled with fluorescent Calcein AM dye) were then incubated on the ECs for 20 minutes to allow binding, washed to remove any unbound U937 cells, and imaged. Image analysis via ImageJ (version 1.54g) showed binding of U937 cells to target PSC-derived ECs, (C) PSC-derived cardiomyocytes (CMs), and (D) PSC-derived cardiac fibroblasts (cFibs) following antibody blocking compared to unblocked controls. **** = p<0.0001, BR=3. Statistical analysis performed via Prism 10.2.2. ROI = Region of Interest.

The advent of CRISPR/Cas9-based gene editing^29,30^ has enabled design of targeted (i.e., cell-specific) gene edits for personalized hPSC-based cellular therapies, potentially overcoming some of the barriers that hampered previous attempts to leverage systemic blockade of ICAM-1 for immune tolerance induction. To assess the potential of ICAM-1 as a hypoimmune target in hPSCs, rather than just in primary cell types as previously described, we first used ICAM-1 blocking antibodies in *in vitro* cultures of clinically-relevant hPSC-based cardiovascular therapies (hPSC-CVTs) to verify inhibition of immune cell binding. We then used a targeted CRISPR/Cas9 editing strategy to create hPSC lines that were null for surface and secreted ICAM-1 expression. ICAM-1 ablation dramatically diminished binding of multiple immune cell types to three different hPSC-CVTs, and impeded *in vitro* and *in vivo* immune responses to the grafts. This new hypoimmune strategy was effective at diminishing immune cell adhesion as a stand-alone gene edit, and when added as a secondary KO to existing first generation hypoimmune MHC KO hPSC lines.

## Results

### ICAM-1 blockade reduces leukocyte adhesion to hPSC-derived cellular therapies

To validate the overall approach of targeting ICAM-1 in hPSCs to promote immune evasion, we first utilized ICAM-1 blocking antibodies for classical *in vitro* binding assays. Multicellular hPSC-CVTs are a promising strategy for generating mature, vascularized grafts to repair damaged heart tissue following myocardial infarction.^31,32^ We focused our *in vitro* studies on three hPSC-derived cell types commonly utilized in these grafts: ECs, CMs, and cardiac fibroblasts (cFibs). ECs were selected as the initial cell type to test the blocking strategy, as they are the first target of allorejection in CVTs and other vascularized hPSC-based therapies, as well as in cardiac allograft vasculopathy and other adverse events that occur during rejection of traditional solid organ transplant grafts.^33,34^ The H9 human embryonic stem cell line was differentiated into highly pure (>90% CD31^+^CD144^+^) ECs using a standardized seven day protocol,^35^ then cultured and stimulated for 48 hours with pro-inflammatory cytokines tumor necrosis factor alpha (TNFα; 10 ng/ml) and interferon gamma (IFNγ; 50 ng/ml) to mimic the early inflammatory post-transplantation microenvironment. Treating H9 ECs with anti-ICAM-1 blocking antibody (0.5 mg/µl) for 1 hour at 37°C resulted in significantly diminished binding of the monocytic U937 cell line to the ECs (Fig. 1b). ICAM-1 blocking also impeded U937 binding in hPSC-derived CMs (Fig. 1c) and cFibs (Fig. 1d). These results implicate ICAM-1 as a critical molecule for immune cell binding to multiple PSC-derived cell types, and are consistent with the literature investigating primary (non-PSC) cell types from humans,^27,36^ mice,^21^ and other species.^37^

### Generation of hypoimmune ICAM-1 KO hPSCs

Having demonstrated the role of ICAM-1 in immune cell binding to hPSC-CVTs and having validated our overall targeting approach, we next employed a CRISPR/Cas9-based strategy to KO ICAM-1 in multiple hPSC lines via introduction of a stop codon, preventing generation of any isoform of the ICAM-1 protein (Fig. 2a). ICAM-1 KO hPSCs were indistinguishable from isogenic wild-type (WT) hPSCs by each of the several commonly-used hPSC characterization metrics we tested. KO lines showed typical hPSC morphology (Fig. 2b). They had similar pluripotency compared to WT, as assessed by staining for SSEA-4 and alkaline phosphatase, and in teratoma assays (Fig. 2c,d and Supplementary Fig. 2a). Cells were karyotypically normal after gene editing as determined by g-banding method (Supplementary Fig. 2b), and no off-target effects of CRISPR/Cas9 editing were detected via Sanger sequencing of the five highest-likelihood off-target sites predicted by CRISPOR algorithms (data not shown). WT and KO lines both displayed normal responses to inflammatory stimulation (TNFα [10 ng/ml] and IFNγ [50 ng/ml] for 48 hours) by upregulating expression of MHC class I on their cell surfaces (Fig. 3a). As expected, WT hPSCs showed baseline expression and upregulation of surface ICAM-1 following inflammatory stimulation, but in the ICAM-1 KO lines no surface protein was detectable by flow cytometry (Fig. 3b). Lastly, we assessed global transcriptional profiles as well as expression of pluripotency-associated genes by bulk RNA sequencing and found no meaningful differences between WT and KO hPSCs (Fig. 3c,d). Our results indicate that our gene editing was effective in ablating functional ICAM-1 protein (i.e., a complete KO) in all lines tested, that the integrity and stability of the hPSC lines is maintained, and that ICAM-1 KO hPSCs are suitable for subsequent differentiation into cellular therapy candidates and immunogenicity studies.

**Figure 2.**
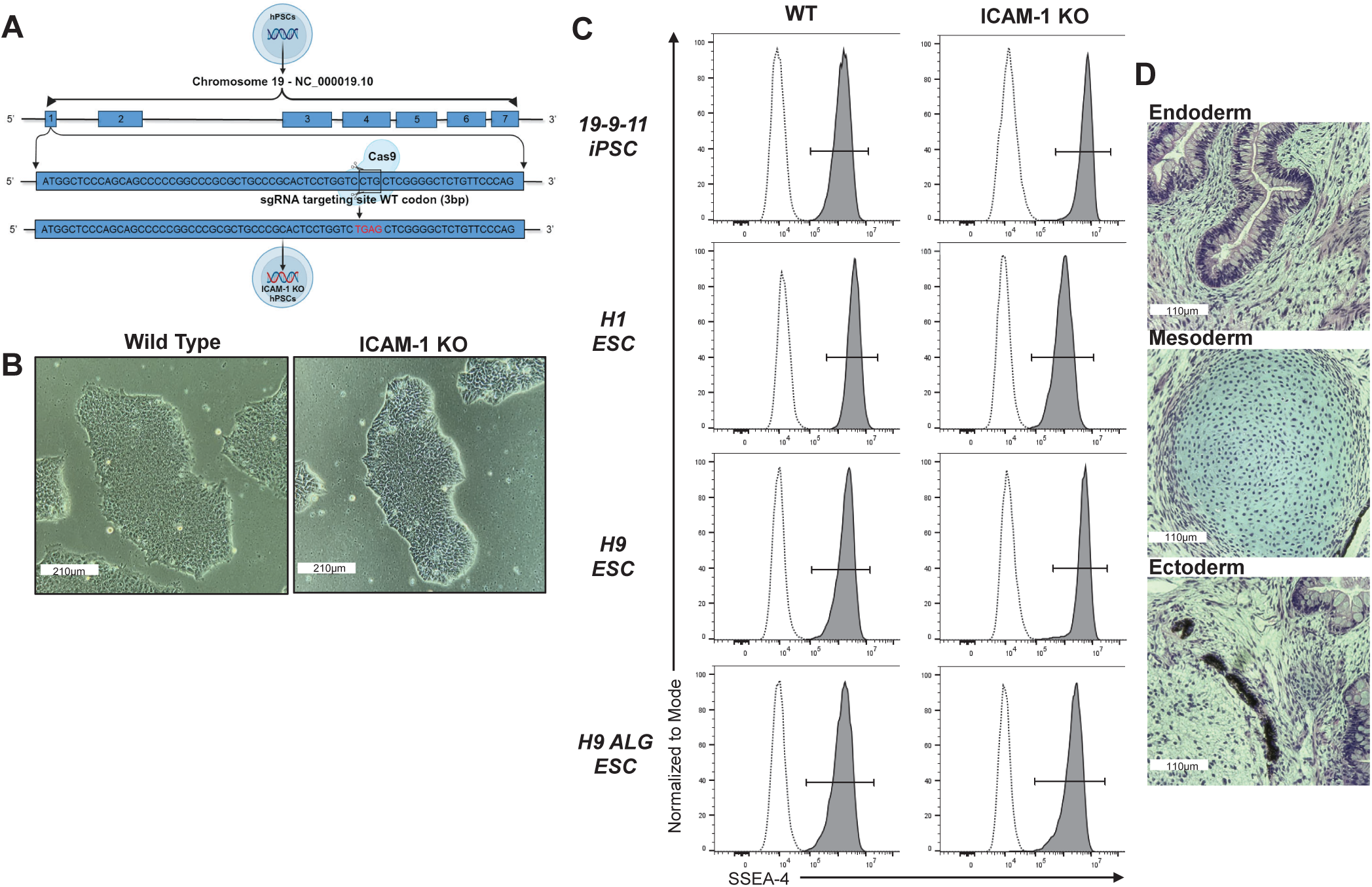
Generation of ICAM-1 Knock-out Pluripotent Stem Cell (PSC) Lines. ICAM-1 was knocked-out (KO) via CRISPR/Cas9 editing. (A) Schematic demonstrating the CRISPR/Cas9 KO strategy. The wild type (WT) codon (CTG) in the first exon of the ICAM-1 sequence is edited into a stop codon (TGA) via the addition of a nucleotide. (B) Representative PSC colony morphology (10X). (C) Pluripotency marker SSEA-4 staining by flow cytometry. (D) Teratomas were grown in immune-deficient mouse hosts by intramuscular injection of the H9 ICAM-1 KO PSC line with Matrigel. Representative Hematoxylin and Eosin stained images of (top) gut [Endoderm], (middle) cartilage [Mesoderm], and (bottom) retinal pigment epithelium [Ectoderm] representing all three germ layers. Multiple cell lines were made in human embryonic stem cells (hESCs) and in human induced PSCs; data are representative of all lines.

**Figure 3.**
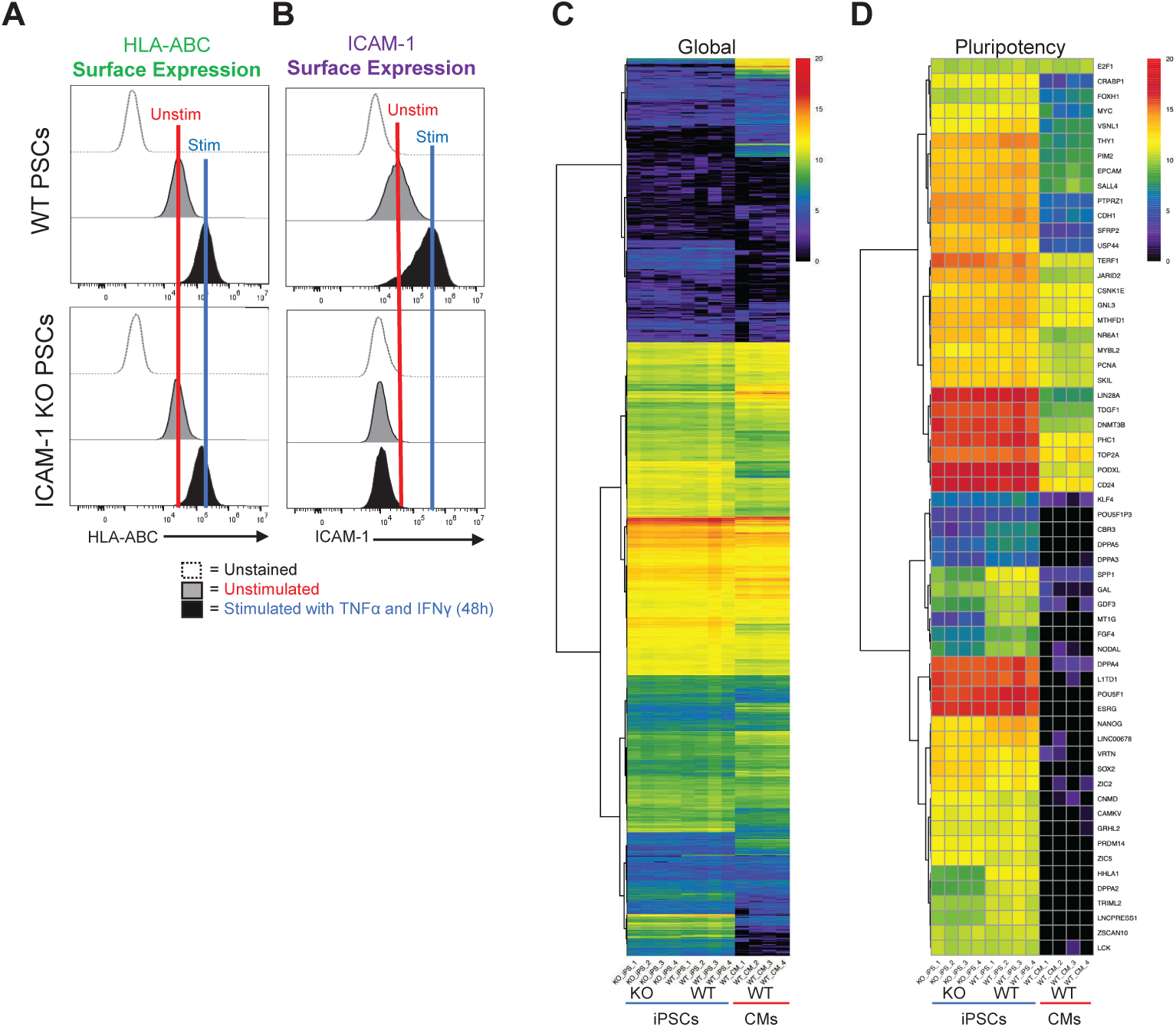
Transcriptomic and Protein Expression Analysis of ICAM-1 Knock-out (KO) vs Wild Type (WT) Cells. ICAM-1 KO and WT isogenic pluripotent stem cells (PSCs) were stimulated with TNFa (10ng/ml) and IFNy (50ng/ml) for 48 hours, then assessed for (A) MHC class I expression and (B) surface ICAM-1 expression at baseline and following stimulation. (C) Bulk RNA sequencing was performed on ICAM-1 KO vs. WT induced PSCs (iPSCs); WT iPSC-derived cardiomyocytes (CMs) shown as a differentiated cell type reference control. Log2 RSEM counts for 21,055 transcripts from four cell culture wells (biological replicates, BRs) of PED05 ICAM-1 KO iPSCs, four BRs of PED05 WT iPSCs, and four BRs of PED05 WT iPSC-derived CMs. Clustering was performed and the graphic was made by running pheatmap in R Studio using Euclidean distance and ward.D hierarchical clustering metrics. (D) Log2 RSEM counts for 60 gene reference markers classifying PSC identity.

### ICAM-1 KO hPSCs have typical differentiation capability

An advantage of hypoimmune gene editing is the theoretical possibility of generating one universal donor hPSC line that can be thoroughly validated for safety and other attributes prior to widespread clinical use in large cohorts of patients. This is in contrast with autologous hPSCs, where each new patient-derived line will likely require batch-specific testing and regulatory approval prior to use.^38^ Given the potential use of hypoimmune hPSCs in multiple disease and clinical contexts, it is critical to assess their capability to differentiate into multiple lineages and validate each universal line’s utility for each disease-specific therapeutic application. As shown in Figure 2, ICAM-1 KO hPSCs formed teratomas and spontaneously differentiated into cell types from each of the three germ layers, indicating robust pluripotency potential. We next assessed the capability of ICAM-1 KO cells to be directed to differentiate into specific hPSC-CVT cell types using well-established protocols.^35,39,40^ Similar to their isogenic WT hPSC controls, ICAM-1 KO cell lines differentiated into high-purity CD31^+^CD144^+^ ECs, cardiac troponin T (cTNT)^+^ CMs, and TE-7^+^ cFibs (Fig. 4a-c). These results show the potential use of ICAM-1 KO hPSCs as a universal cell line for generation of reparative CVTs. We next needed to assess immune cell interactions, as this metric is critical for determining their full clinical potential.

**Figure 4.**
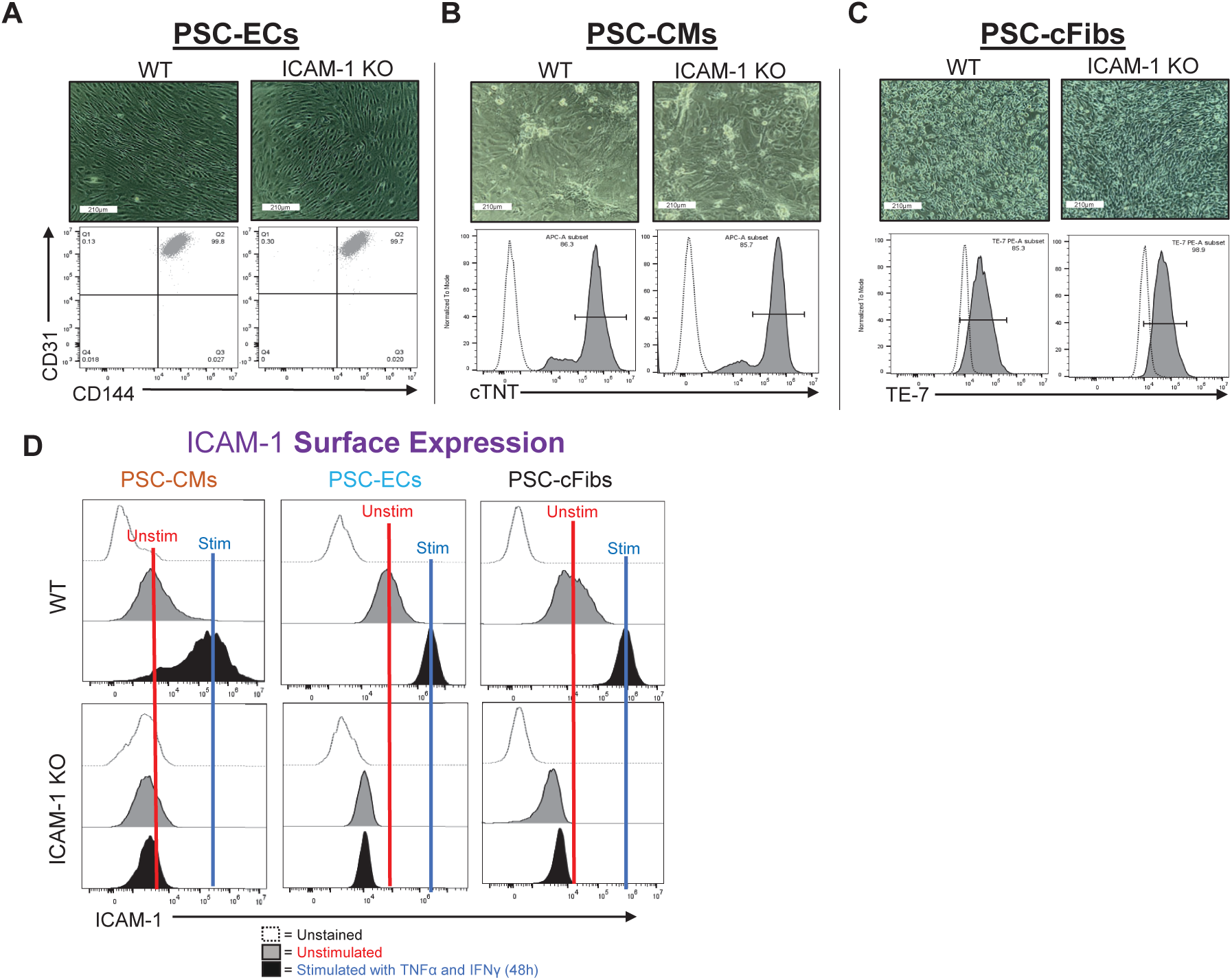
ICAM-1 Knock-out (KO) Pluripotent Stem Cell (PSC) Differentiation Capacity and Surface ICAM-1 Protein Expression. Morphology of (A) PSC-derived cardiomyocytes (CMs), (B) PSCderived endothelial cells (ECs), and (C) PSC-derived cardiac fibroblasts (cFibs) (top panels) is shown (10x). The purity of cell-type specific markers was determined by flow cytometry (bottom panels). (D) ICAM-1 surface expression is shown at unstimulated baseline and following 48 hours of stimulation with TNFα (10ng/ml) and IFNγ (50ng/ml). Data are representative of n=3 experiments, with triplicated biological replicates. Analysis was performed via FlowJo 10.10 software. Other abbreviations: WT – wild type.

### Surface and soluble ICAM-1 expression is ablated on differentiated cells

In humans and mice, there are multiple isoforms of ICAM-1 generated via alternative splicing and proteolytic cleavage events that occur during inflammatory responses. These isoforms are expressed at the cell surface and in a secreted soluble form (sICAM-1), so we assessed our hypoimmune hPSCs for both. Similar to the parent hPSCs, there was no surface ICAM-1 detected in any of the hPSC-differentiated cell types by flow cytometry (Fig. 4d), indicating that the KO was thorough and effective, devoid of the leakiness that has been documented in some previous hypoimmune gene editing approaches (e.g., MHC class I via KO of beta-2 microglobulin [β 2M]).^10,41^ The differentiated cells also maintained normal MHC class I upregulation responses to inflammatory stimuli (Supplementary Fig. 3).

Interestingly, in the late 1990s/early 2000s researchers did not yet know of ICAM-1 alternative splicing and multiple groups generated ICAM-1-deficient mouse strains that were not completely null for all functional isoforms. Indeed, as described by Ramos et al., the majority of all existing literature describing ICAM-1 KO mice does not include true ICAM-1 KOs, but rather mice only deficient in specific isoforms.^42^ While ICAM-1 KO mice have since been generated,^43^ the effect of this significant oversight still impacts the knowledge base in this area. Our gene editing strategy introduces a stop codon in the first exon of the ICAM-1 gene on Chromosome 19, preventing translation and subsequent formation of any complete isoform protein. No surface protein is detectable in any of the n=6 independently-generated ICAM-1 KO hPSC lines made by our team. To validate KO of sICAM-1 (i.e., complete nullification of all ICAM-1 protein products), we performed a Luminex assay on cell culture supernatants from hPSC-ECs that had been stimulated, as described in Figures 1, 2, and 4 above, for 48 hours with inflammatory cytokines. WT hPSC-ECs secreted large quantities (>100 nanograms/ml) into cell culture supernatants following stimulation, while KO cells have no detectable levels (Fig. 5). These results confirm the effectiveness of our KO strategy for total ablation of *surface* as well as *secreted* isoforms of ICAM-1. As sICAM-1 has been used as a biomarker for a number of inflammation-associated pathologies, including spontaneous and transplantation-associated coronary artery disease,^44^ the impact of its absence in future hypoimmune hPSC therapies may have relevance for tissue repair following transplantation in terms of both biological function and clinical assessment of graft failure.

**Figure 5.**
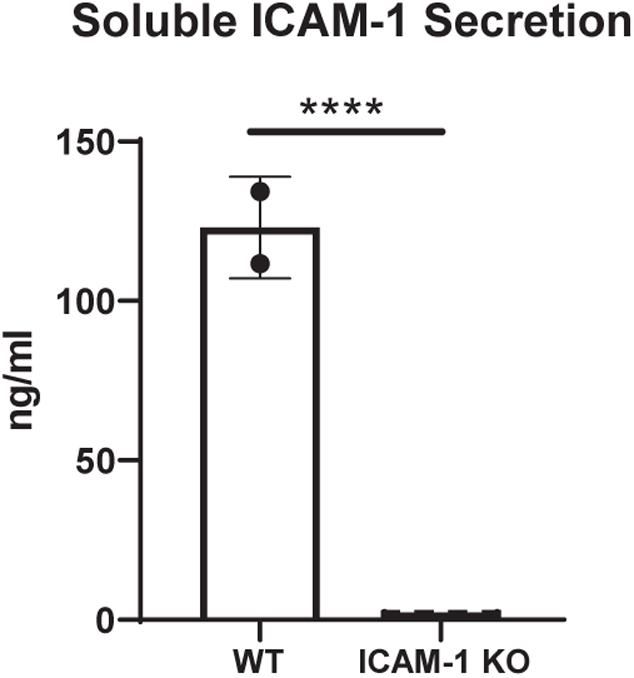
Soluble ICAM-1 Secretion. Pluripotent stem cells were differentiated into highly pure (>96% CD31+CD144+) endothelial cells and stimulated with 10 ng/ml TNFα and 50 ng/ml IFNγ for 48 hours. Soluble ICAM-1 secreted into the cell culture media was assessed by Luminex assay. Unpaired t test, **** = p<0.0001, n=2 biological replicates for wild type (WT) and n=4 biological replicates, n=3 independent experiments. Statistical analysis was performed via Prism 10.2.2 software. Data are representative of n=3 experiments. Other abbreviations: KO – knock-out.

### Immune cell adhesion is diminished in ICAM-1 KO cells

Immune cell binding to ECs (as well as to CMs, cFibs, and other parenchymal cells in various other transplanted grafts) is stabilized via ICAM-1 and is a critical first process in the early, inflammatory stages of allograft rejection following solid organ and cellular therapy transplantation. These interactions continue to imperil transplanted grafts during the temporal processes of acute and chronic allorejection by driving the formation of immune cell synapses required for effector immune cell-mediated cytotoxicity events. A central premise of our study is that impeding this key first step of allorejection—immune cell binding—will diminish downstream pathologic immune responses and therefore improve post-transplantation hPSC graft survival.

Monocytes and macrophages are integral components of the innate immune system that play crucial roles in the rejection of transplanted tissues. Following transplantation, these cells are swiftly recruited to the graft site where they serve as central mediators of inflammatory responses.^45^ Monocytes differentiate into macrophages following ICAM-1-mediated extravasation into the graft tissue, where they engage in phagocytosis of foreign material and initiate immune surveillance. Through the secretion of pro-inflammatory cytokines and chemokines, as well as antigen presentation to T cells, macrophages contribute to the activation and recruitment of other immune cells, ultimately promoting graft rejection. Moreover, macrophages are implicated in chronic rejection processes, such as fibrosis and tissue remodeling, which can lead to long-term graft dysfunction. Overall, the dynamic interplay between monocytes and macrophages underscores their pivotal role in orchestrating immune responses against transplanted organs, highlighting them as potential targets for therapeutic intervention aimed at prolonging graft survival. To assess this immune cell binding, we first used the well-established U937 monocytic lymphoma cell line, which expresses the ICAM-1 binding ligands LFA-1 and MAC-1 (see Fig. 1a), to model early innate immune cell interactions with hPSC-ECs following transplantation. High purity hPSC-ECs were differentiated from WT and ICAM-1 KO lines, stimulated for 48 hours with TNFα (10 ng/ml) and IFNγ (50 ng/ml), then co-cultured as described in Figure 1 with fluorescently-labeled U937 cells. ICAM-1 KO ECs showed dramatically diminished binding of U937s compared to WT controls (Fig. 6).

**Figure 6.**
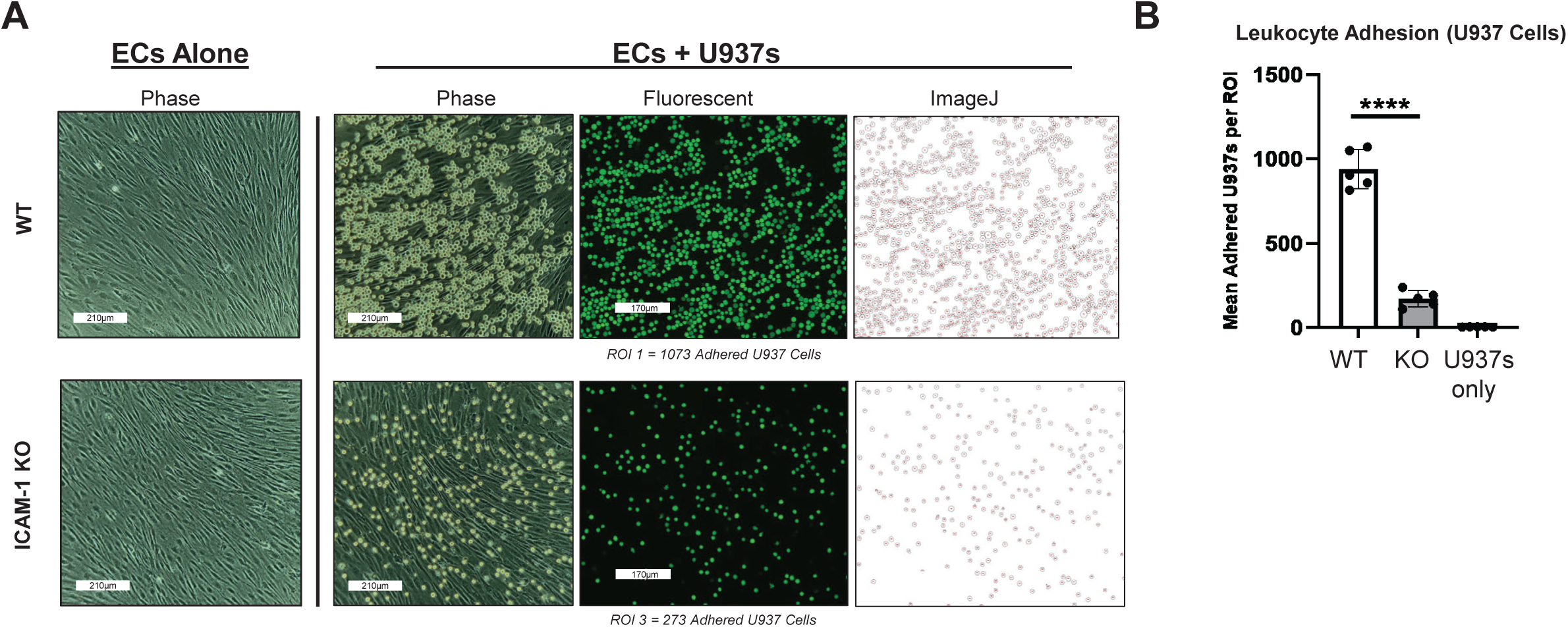
Immune Cell Adhesion Assay with ICAM-1 Knock-Out (KO) Pluripotent Stem Cell (PSC)-Derived Endothelial Cells (ECs). H9 wild type (WT) and ICAM-1 KO PSCs were differentiated into high-purity CD31+CD144+ ECs and stimulated with TNFα (10ng/ml) and IFNγ (50ng/ml) for 48 hours. As in Figure 1, U937s were co-cultured with both cell types and washed. Images were acquired on ECHO Revolve | R4 microscope. **** p<0.0001, biological replicates=3. Statistical analysis was performed via Prism 10.2.2 software. ROI = Region of Interest

Neutrophils are another primary responder of the innate immune system and they also play a significant role in the rejection of solid organ transplants. Upon transplantation, neutrophils are rapidly recruited to the site of graft placement where they unleash a cascade of pro-inflammatory mediators and engage in phagocytosis to eliminate foreign antigens.^46^ Although traditionally considered short-lived effector cells, recent studies have highlighted their involvement in orchestrating sustained inflammatory responses and modulating adaptive immunity.^47^ Neutrophils contribute to tissue damage through the release of reactive oxygen species, proteases, and pro-inflammatory cytokines, exacerbating graft injury and promoting immune activation. Furthermore, their interactions with other immune cells, such as dendritic cells and T cells, further amplify the alloimmune response and contribute to the pathogenesis of graft rejection.^45^ Thus, understanding the multifaceted role of neutrophils in transplant rejection and experimentally testing the neutrophil binding response to hPSC-derived grafts is crucial for mitigating their detrimental effects and prolonging graft survival.

Similar to the U937 binding experiments described above, we fluorescently labeled allogeneic neutrophils and co-cultured them with hPSC-ECs, showing diminished binding to ICAM-1 KO cells (Supplementary Fig. 4). We also performed binding experiments with peripheral blood mononuclear cells (PBMCs; i.e., a mixture of innate and adaptive LFA-1 and MAC-1 positive cells, but lacking neutrophils or other granulocytes) from n=2 human leukocyte antigen (HLA)-mismatched donors (Supplemental Fig. 1a-c, 5, Table 1). All innate and adaptive immune cells tested in this series of experiments showed significantly diminished binding to hPSC-derived cells, indicating a broad anti-adhesion effect of ICAM-1 KO.

### ICAM-1 KO is protective against *in vitro* and *in vivo* immune responses

ICAM-1 KO, similar to antibody blocking, dramatically reduces but does not completely prevent binding of innate and adaptive immune cells to hPSC-derived cells. We hypothesized that the very few numbers of immune cells that do attach to hPSC-ECs would have attenuated immune responses to the other ICAM-1 KO cell types within the hPSC-CVT. We conducted mixed lymphocyte reactions to assess the alloantigen-induced proliferative responses of HLA-mismatched T cells to the hPSC-CVTs (i.e., to model the effect of cells that bound ECs and were able to extravasate into the graft, then encountering ICAM-1 KO parenchymal cells). In response to 6 day mixed lymphocyte reaction co-culture with hPSC-CMs, we observed significantly diminished proliferative responses of total CD8+ T cells, as well as the CD8+ effector memory (T_EM_) CD45RO^+^ CD45RA^-^ CD62L^-^ CCR7^-^ cell subset known to play a prominent role in T cell-mediated rejection^48^ (Fig. 7a,b). These data indicate that the lack of ICAM-1 binding resulted in a diminished proliferation response in this alloreactive T cell population.

**Figure 7.**
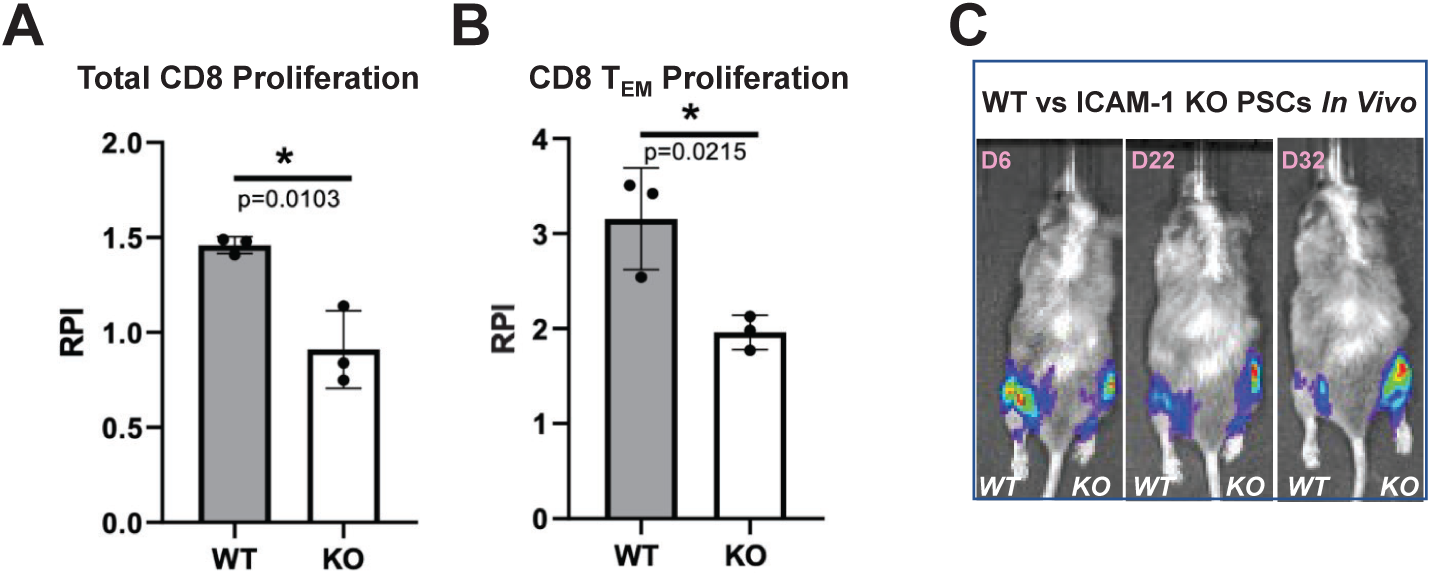
In Vitro and In Vivo Immune Responses to ICAM-1 Knock-out (KO) Cells. To test in vitro immune responses to edited cells, highly pure (>85% cTNT+) PSC-derived CMs were made from wild type (WT) and ICAM-1 KO pluripotent stem cells (PSCs), similar to Figure 1C. Cells were co-cultured for 6 days with HLA-mismatched peripheral blood mononuclear cells (1:6 T:E ratio) labeled with VPD450 proliferation dye in a mixed lymphocyte reaction. Alloreactivity was assessed via proliferation (VPD450 dye dilution) of total viable cells and various T cell subpopulations by flow cytometry. Proliferation of (A) Total CD8+CD3^+^ T cells and (B) Effector memory T (TEM) cells (CD45RO+ CD45RA-CD62L-CCR7-) is shown. Relative proliferation index (RPI) is the ratio of % proliferating cells with cardiomyocyte targets/baseline proliferating cells without targets. N=3 biological replicates, error bars = standard deviation, analysis via 2-tailed unpaired t-test, Prism 9.3.1. N=3 repeat experiments. (C) To test in vivo immune responses to edited cells, NeoThy humanized mice with flow-confirmed hCD45+ and hCD3^+^ (both >10%) immune cells were engrafted with 1e6 ICAM-1 KO (right leg) vs. WT (left leg) isogeneic PSCs co-injected with Matrigel. Humanized immune systems were HLA-mismatched to the PSC grafts. Bioluminescence imaging (BLI) signal was monitored for 32 days at early (6 days), middle (22 days), and late/terminal (32 days) time points. A representative mouse is shown reflecting loss of WT graft, and retention of KO graft, which was seen in 4 of 5 mice BLI signal quantified by total flux (p/s).

We then tested *in vivo* immune responses to hypoimmune cells using the NeoThy, an advanced human immune system humanized mouse model created by intravenous injection of neonatal human cord blood hematopoietic stem and progenitors and surgical implantation of human thymus tissues.^49,50^ Similar to the fetal tissue-based bone marrow-liver-thymus (“BLT”) mouse model, NeoThy mice engraft with a robust repertoire of adaptive and innate immune cells but use more developmentally-mature non-fetal tissue sources. We transplanted WT and ICAM-1 KO hPSCs expressing constitutive Akaluc bioluminescence imaging (BLI) reporter constructs, co-injected with Matrigel, into the hind limbs of NeoThy mice with verified human immune chimerism (made with HLA-mismatched humanizing tissues) exceeding 10% human CD45^+^ and 10% human CD3^+^ cells in circulation. In each individual humanized recipient (n=5), WT cells were injected into the left mouse leg and KO cells into the right leg (Fig. 7c). In 4 of 5 mice, KO cells were retained and grew into teratomas, similar to non-humanized controls used for the teratoma assays in Figure 2d. In contrast, WT cells were controlled or diminished in size, as evidenced by live animal imaging of BLI signal over three timepoints up to 32 days post-transplant. Taken together, the *in vitro* and *in vivo* immune assay results indicate an overall differential and more robust immune response to WT cells compared to KO cells. The previous binding assay data offer additional evidence implicating immune cell binding, or lack thereof, as a critical mechanism mediating these results. Overall, these findings support the use of ICAM-1 KO hPSCs as a hypoimmune therapy platform with potential to elicit attenuated, or potentially to completely evade, allorejection responses following transplantation into patients.

### ICAM-1 KO confers additional protection to first-generation hypoimmune hPSCs

There are a variety of promising hypoimmune gene editing strategies to prevent adaptive immune responses to hPSC-derived grafts,^7,9,10,51^ but to date no strategy has been demonstrated to blunt all components (cellular and humoral) of the multifaceted allorejection response. Similar to xenotransplantation gene editing strategies,^52^ it may be useful to KO multiple genes in a single universal hypoimmune hPSC line to promote evasion of the coordinated multicellular immune responses that occur *in vivo*. Such strategies will need to be validated in multiple hPSC-based cell therapies and in several disease contexts, and certain disease-specific therapies may need more editing than others. Additionally, introduction of multiple gene edits in one cell line will need to balance effectiveness with avoidance of off-target effects and other adverse events.^53,54^

Having demonstrated that ICAM-1 KO diminishes immune cell binding as a stand-alone edit, we sought to validate addition of ICAM-1 KO to a promising, previously published first generation hypoimmune hPSC line to test whether it was possible to confer anti-adhesion properties to the line. We first engineered the H9 human embryonic stem cell PSC line with a constitutively expressed Akaluc GFP construct to enable BLI,^55^ followed by a double (β 2M and CIITA) KO coupled with overexpression of natural killer cell inhibitory ligand CD47, as described by Deuse et al.^9^ This line was called “H9 DKO.” We then added ICAM-1 KO to H9 DKO, as described for editing of WT lines above, to make the “H9 TKO” line. As expected, after differentiation into highly pure ECs, both H9 DKO and H9 TKO did not express or upregulate MHC class I after stimulation for 48 hours with TNFα (10 ng/ml) and IFNγ (50 ng/ml). Cell surface ICAM-1 and sICAM-1 expression at baseline and following stimulation was still present in H9 DKO, indicating that ICAM-1-mediated immune cell binding and associated allorejection events are still an active concern for first generation hypoimmune hPSCs. As expected, ICAM-1 KO ablated all isoforms of ICAM-1 in H9 TKO (Fig. 8a,b). Concordantly, the ICAM-1 null H9 TKO line elicited diminished binding of U937 cells (Fig. 8c) and PBMCs (Fig. 8d) following stimulation in leukocyte adhesion assays. These results are important proof-of-concept data showing additive benefits of introducing ICAM-1 KO into existing hypoimmune gene edited hPSC lines, which could be beneficial for creation of optimized universal donor hPSC cells for specific pathological contexts. These experiments did not directly compare the overall immune evasion capability of single ICAM-1 KO edited hPSC lines vs the H9 TKO line, as this was outside the scope of this study.

**Figure 8.**
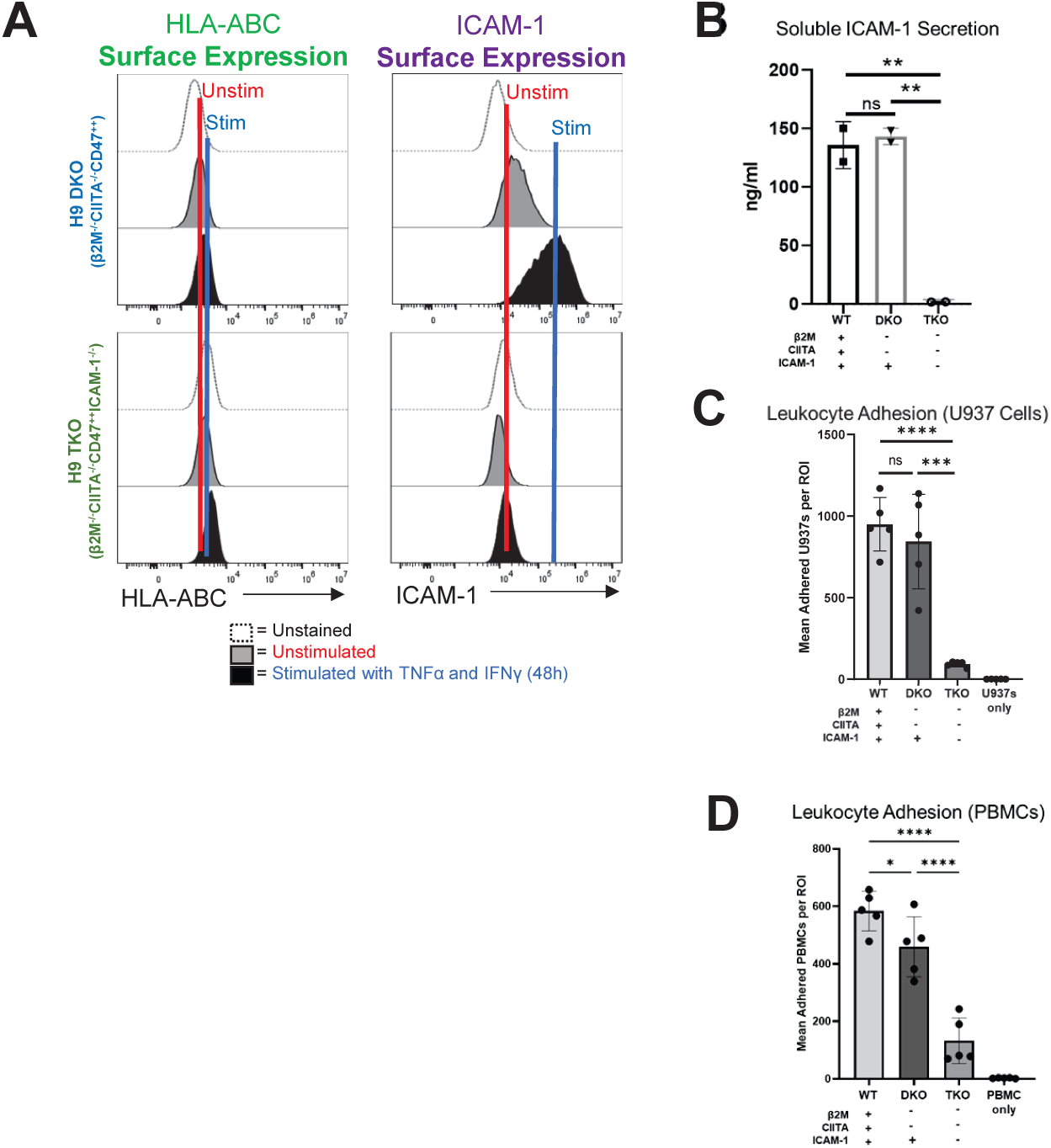
Addition of ICAM-1 Knock-out (KO) to First Generation Hypoimmune Pluripotent Stem Cell (PSCs). (A) “First generation” B2M−/−, CIITA−/−, CD47++ (DKO) PSCs were edited to introduce the ICAM-1-KO, making a triple KO (TKO) cell line. PSCs were differentiated into highly pure (>98% CD31+CD144+) endothelial cells (ECs) and stimulated with 10 ng/ml TNFa and 50 ng/ml IFNy for 48 hours and assessed for HLA class I (left) and ICAM-1 (right) surface protein expression. (B) Wild type (WT), DKO, and TKO PSCs were differentiated into highly pure (>98% CD31+CD144+) ECs and stimulated as above, then assessed by Luminex for secreted soluble ICAM-1, as in Figure 5. (C) Similar to Figures 1 and 6, adhesion assays were performed using U937s as immune effector cells. U937s were stained with violet VPD450 dye. (D) Adhesion assays were also conducted with peripheral blood mononuclear cells (PBMCs). N=3 biological replicates, error bars = standard deviation, analysis via 2-tailed unpaired t-test. WT = wild-type H9 PSC line with ALG (Akaluc + GFP) reporter construct, DKO = H9 PSC line with KO of β2M and CIITA, with ALG reporter construct, TKO = DKO line with addition of ICAM-1 KO. * p<0.05, ** p<0.01, *** p=0.0001, **** p<0.0001, ns = not significant. Statistical analysis was performed via Prism 10.2.2 software.

## Discussion

A robust allorejection response can swiftly destroy even a functionally perfect cellular therapy. It is therefore imperative to develop effective strategies to enable hPSC therapies to engraft and survive encounters with a transplant recipient’s immune system. Off-the-shelf hypoimmune edited hPSCs have tremendous promise in this regard, but existing gene editing strategies have not fully addressed important immune subsets that participate in early graft rejection. Our study introduces a novel approach to hPSC hypoimmune therapy design by directly targeting ICAM-1, a common adhesion molecule involved in multiple aspects of both the adaptive and innate immune responses to transplanted grafts.

The strategy of ablating/blocking ICAM-1 to diminish effector immune responses was explored prior to the advent of hypoimmune hPSC gene editing, but a number of early studies relied upon flawed mouse models that harbored incompletely KO’d ICAM-1 isoforms, potentially skewing aspects of the canon of knowledge. Similarly, early recombinant antibody and antisense oligonucleotide therapies targeting ICAM-1 have been assessed in transplantation and other contexts (e.g., intestinal inflammation),^26^ but as with other recombinant antibody-based therapeutic approaches, design considerations^56,57^ and the systematic nature of these treatments (i.e., binding of ICAM-1 on target ECs as well as to off-target immune and parenchymal cells) may not have the advantages that precise cell-type specific gene editing can offer. In any event, it may be prudent to revisit the early literature and repeat selected experiments using the more advanced tools that are now at our disposal.

In this study, we leveraged two critical technological advances (hPSCs and CRISPR/Cas9 gene editing) to precisely introduce ICAM-1 KO into hPSC-CVT cell types. We showed that ICAM-1 KO has clear effects in diminishing binding of innate and adaptive immune cells and that evasion of *in vitro* and *in vivo* immune recognition is conferred by this editing approach. We also demonstrated that addition of ICAM-1 KO to first generation β 2M and CIITA double KO/CD47 overexpressing cells results in karyotypically normal hPSCs with diminished immune cell-binding propensity, in addition to retaining the original hypoimmune benefits of the parent line. Future studies will be needed to discern whether stand-alone ICAM-1 KO is sufficient for all clinical applications or whether combining ICAM-1 KO with other strategies is ideal. ICAM-1 is known to participate in T cell activation, and our *in vitro* T cell proliferation study showed that ICAM-1 KO directly impacts T cell alloresponses; hypoimmune hPSCs may therefore not need MHC KO to mitigate T cell-mediated rejection. Antibody-mediated rejection, however, is an important driver of allorejection and MHC I and II are primary targets for donor-specific antibodies, so MHC KO combined with ICAM-1 KO may still be advantageous for transplanting into highly-sensitized patients.^58^ Careful crossmatch assessment prior to transplantation could potentially minimize the incidence of antibody-mediated rejection in hPSC-CVT transplantation.^59,60^ It should be noted that antibody-mediated rejection can involve antibodies targeting a variety of non-HLA epitopes (e.g., EC-specific surface molecules),^61–63^ so it may not be possible to create a universal cell line via MHC ablation alone that is completely effective in eliminating this powerful driver of allorejection. Other additive hypoimmune editing approaches may be needed, such as a promising CD64 overexpression strategy recently described by Gravina et al.^51^ Relatedly, better *in vitro* and *in vivo* models of human antibody-mediated rejection are needed to adequately interrogate these types of research questions. Humanized mouse models are quite useful for T cell-mediated rejection and natural killer cell-mediated allorejection studies,^64–67^ but the most commonly used humanized mice are suboptimal for antibody-mediated rejection studies due to B cell immaturity and impeded class switching in *de novo* antibodies.^68^ Our group and others are working on new iterations of humanized mouse models that incorporate more robust T cell, natural killer cell, and innate immune cell repertoires in order to improve their utility for transplantation immunology-focused research studies, and creation of tractable models harboring antigen-specific IgG antibody pools is an area of intense focus. We anticipate that improved next-generation humanized mouse models will become a critical tool in the clinical translation of new hypoimmune gene edited hPSC therapies in the coming years.

Future applications of ICAM-1 KO and related editing could include optimization of engineering strategies to attenuate, but not totally ablate, immune responses in order to balance diminishing allorejection and graft loss with allowing for some amount of residual immunosurveillance capacity against malignant transformation and/or formation of viral (e.g., cytomegalovirus) reservoirs. Additionally, our ICAM-1 KO developed for use in hPSC therapeutics could also be a promising target for gene editing in xenotransplantation. ICAM-1 is highly conserved across species, and previous studies have shown that human cells bind to pig cells via ICAM-1.^69^ Lastly, ICAM-1 is the primary receptor for human rhinovirus,^70^ and also plays a role in cell-to-cell transmission of human immunodeficiency virus.^71^ We intend to explore the impact of ICAM-1 KO on viral infection dynamics in future studies.

Our study highlights the importance of designing hypoimmune gene editing approaches with adaptive and innate immune cells in mind. Targeting ICAM-1 as a common molecule utilized by multiple cell types during the earliest steps in the allorejection process demonstrates a new paradigm for developing novel hypoimmune gene edits. Further progress in this area of research will ultimately improve the long-term efficacy and safety of regenerative therapies for cardiovascular pathologies and a number of other diseases.

## Materials and Methods

This work was approved by the Animal Care and Use Committee of the University of Wisconsin (UW) School of Medicine and Public Health and the UW Health Sciences Institutional Review Board, and complied with federal and state law.

### Genetic Modification of PSC Lines

#### KO of ICAM-1

Multiple human induced PSC and embryonic stem cell lines were edited. The single guide RNA (sgRNA) for the site of interest was identified using the CRISPOR design tool.^72^ 1.5 nmol synthetic sgRNAs were constructed with 2’-O-methyl 3’ phosphorothioate modifications at the first and last 3 nucleotides (Synthego). In preparation for electroporation, PSCs were cultured on Matrigel in TeSR-PLUS media (StemCell Technologies) and grown to ∼80% confluency following standard cell culture protocols. Cells were treated with CloneR (StemCell Technologies) 24 hours before electroporation per the manufacturer’s protocol. Preceding electroporation, reconstitution of sgRNA constructs to a concentration of 150 pmol/mL was conducted following the manufacturer’s protocols. We combined 4 mg of Cas9 Nuclease protein (TrueCut Cas9 Protein V2, Thermo Fisher Scientific), 5 mL of Neon Buffer R (Invitrogen), and 1 mL of reconstituted sgRNA to promote Cas9-ribonucleoprotein (RNP) complex formation. After an incubation of 15 minutes, 1.0 mL of ssODN primer designed with homology overhangs of at least 40 base pairs (reconstituted to 1 mg/mL) was added into the Cas9-RNP mixture. PSCs were detached and singularized utilizing a 1:1 mixture of 0.5 mM EDTA and Accutase (Invitrogen) via incubation with EDTA/Accutase for 3-4 minutes, resuspension in 1 mL phosphate buffered saline (PBS; Corning), and centrifugation into a pellet. The cells were resuspended in 35 mL Buffer R (Thermo Fisher Scientific) and combined with 8 mL of the previously prepared Cas9-RNP complex with repair ssODN. Cells underwent electroporation via 10 μL NEON electroporation format with 1x pulse, 30 msec, and 1200V settings. Following four subsequent rounds of electroporation, cells were pooled and plated in TeSR-PLUS (StemCell Technologies) maintenance media supplemented with CloneR (StemCell Technologies) per the manufacturer-recommended concentrations utilizing serial dilutions to foster single-cell clonal growth. After expansion over a span of 10-14 days, clones were identified and picked using standard techniques. QuickExtract DNA Extraction Solution 1.0 (Epicentre) was used to generate bulk gDNA from dissociated cells to validate editing efficiency before clonal selection. Single-cell clones were manually selected and subsequently mechanically disaggregated. gDNA isolation was performed from a portion of the clones via QuickExtract DNA Extraction Solution 1.0 (Epicentre). Genotyping primers were created by designing flanks around the mutation site, promoting amplification of this region via Q5 polymerase-based PCR (New England Biolabs). PCR products were then identified by agarose gel method and purified via Zymoclean Gel DNA Recovery Kit (Zymo Research). Sanger sequencing of clones was performed by Quintara Biosciences to identify clones that displayed the proper genetic modification. Suspected off-target sites for genome modification were analyzed to identify if the CRISPR/Cas9 system had generated any non-specific genome editing. Genotyping primers were designed to amplify the 5 highest-likelihood off-target sites for each sgRNA as predicted by the CRISPOR algorithms. The resulting PCR products were identified by agarose gel method, purified via Zymoclean Gel DNA Recovery Kit (Zymo Research), and submitted for Sanger sequencing to Quintara Biosciences.

#### Knock-In of Constitutive Reporter Constructs

Generation of base PSC lines with BLI and fluorescence reporters for downstream studies and additional editing (detailed below) was conducted as described in Zhang et al.^55^ The H9 (WA09) embryonic stem cell line (WiCell) and the PED05 induced PSC line^50^ were edited for constitutive expression of the Akaluc bioluminescence reporter as well as green fluorescence protein (GFP) from Aequorea victoria.^73^ The 5’– and 3’-homology arms of the targeting vector were cloned into a drug resistance containing vector. gRNA was synthesized according to the manufacturer’s instructions (GeneArt™ Precision gRNA Synthesis Kit, Thermo Fisher Scientific). To achieve the best electroporation efficiency, hPSCs were passaged with EDTA (1:4 split) and cultured to reach 80–90% confluency 2 days before the experiment. On the day of the experiment, hPSCs were resuspended at a density of 1.25 x 10^7^ cells/mL in E8 medium supplemented with 10 μM Y27632. To prepare RNP, 0.6 µg gRNA was mixed with 2 µg Cas9 protein (TrueCut™ Cas9 Protein v2, Thermo Fisher Scientific) in 5 µl E8 medium supplemented with 10 μM Y27632. The mixture was incubated for 10 minutes at room temperature. Next, we added 5.5 µl cell suspension and 4 μg targeting vector to RNP. The electroporation was performed in the Neon Electroporation System (Thermo Fisher Scientific) with program 13. Cells were combined from 2-4 electroporations and seeded into 1 10-cm dish coated with Matrigel. Drug selection was performed when cells reached 20% confluency. Surviving colonies were picked after drug selection and expanded in E8 medium. The edited lines were called H9 ALG and PED05 ALG, respectively, and used as WT (non-hypoimmune) controls for BLI studies.

#### Generation of Human MHC Double KO-CD47 Overexpression Cell Lines

The H9 ALG line was also then edited using the Deuse et al. strategy^9^ to overexpress CD47 (natural killer cell inhibitory molecule), in addition to KO ofβ 2M (to prevent MHC I surface expression) and CIITA (to prevent HLA class II gene expression and thus inhibit MHC II surface expression).β 2M and CIITA gRNA were synthesized according to the manufacturer’s instructions (GeneArt™ Precision gRNA Synthesis Kit, Thermo Fisher Scientific). H9 ALG cells were resuspended at a density of 1.25 x 10^7^ cells/mL in E8 medium supplemented with 10 μM Y27632. To prepare RNP, 0.3 µg of each gRNA was mixed with 2 µg Cas9 protein (TrueCut™ Cas9 Protein v2, Thermo Fisher Scientific) in 5 µl E8 medium supplemented with 10 μM Y27632. The mixture was incubated for 10 minutes at room temperature. Next, a 5.5 µl cell suspension was added to RNP, and electroporation was performed in the Neon Electroporation System (Thermo Fisher Scientific) with program 16. To get single-cell clones without drug selection, 2.5% of the electroporated cells were seeded into 1 10-cm dish coated with Matrigel. Single cell-derived clones were picked 1-2 weeks later without drug selection. This first generation hypoimmune line with bioluminescence reporter is referred to as H9 DKO. The H9 DKO line was further edited to KO ICAM-1 expression, as described above. The resulting line is referred to as H9 TKO.

### Karyotyping

G-banding analysis was conducted by the WiCell Research Institute (Madison, WI) in accordance to a standardized protocol.^74^ Cultures were harvested following standard metaphase preparation procedures. Briefly, cultured cells were exposed to ethidium bromide as an intercalation agent followed by Colcemid (Thermo Fisher Scientific), which arrests the cells in metaphase. Once the incubation is complete, the cells are exposed to potassium chloride followed by Carnoy’s fixative. Fixed cells are dropped onto slides in an environmental chamber, banded with trypsin, and stained with Leischman’s stain before imaging and analysis.

### SSEA-4 Flow Cytometry

PSCs cultured on Matrigel-coated (Corning) cell culture plates were harvested and stained for the presence of SSEA-4 (clone MC813-70, Thermo Fisher Scientific) to characterize pluripotency per the manufacturer’s instructions. Each well was aspirated of residual cell culture media and incubated for 7 minutes at 37°C with 1 mL 1X TrypLE (Gibco) to detach the cells from the plate. After incubation, 1 mL of MACs Separation Buffer (Miltenyi) was added to each well as a staining buffer to collect and transfer the detached cells to a microcentrifuge tube. The cells were then centrifuged at 300 x g for 5 minutes. The supernatant was aspirated, and each tube was stained with 5 μL of SSEA4 antibody and incubated in the dark for 20 minutes at 4°C. After the incubation, the cells were centrifuged at 300 x g for 5 minutes and the supernatant was aspirated. Flow cytometry was performed on a Cytoflex Flow Cytometer (Beckman Coulter) and analyzed using FlowJo Version 10.9.0 software.

### Alkaline Phosphatase Assay

All PSC lines were grown to near-confluence on Matrigel-coated (Corning) cell culture plates. Cells were then stained with Vector Blue Alkaline Phosphatase Substrate Kit III (Vector Laboratories) according to the manufacturer’s protocol to assess pluripotency.

### PSC Culture

All PSC lines were cultured on Matrigel-coated (Corning) cell culture plates and fed daily with 2.5 mL TeSR E8 media (StemCell Technologies, WiCell Formulation). Upon reaching full confluency, cells were passaged to fresh Matrigel-coated cell culture plates. Each well was aspirated of residual cell culture media and rinsed with 2 mL PBS (without Calcium or Magnesium [−/−]; Corning). To detach cells from the plate for passaging, each well was incubated with 1 mL 5mM EDTA solution made by adding 500μL 0.5M EDTA (Thermo Fisher Scientific) to 500 mL PBS−/− at room temperature for 7 minutes. After incubation, each well was aspirated and TeSR E8 media (StemCell Technologies, WiCell Formulation) was used to dislodge the remaining colonies still attached to the wells and transfer the cells into a fresh Matrigel-coated cell culture plate. Passage number was recorded, with each new passage adding to the cumulative total (i.e., passage number was not reset upon gene editing but continued onward from the input passage number at the start of the editing process).

### Differentiation of Cardiovascular Cells from PSCs

#### EC Differentiation

PSC-derived ECs were differentiated from all the PSC lines using the STEMdiff Endothelial Differentiation Kit (StemCell Technologies) according to the manufacturer’s protocol. Prior to differentiation, PSCs were maintained in TeSR E8 media on Matrigel-coated cell culture plates until they reached full confluency. Following 7 days of differentiation, ECs cultured on Matrigel-coated cell culture plates were aspirated of residual Endothelial Induction Media (StemCell Technologies) and incubated for 5 minutes at 37°C with 1 mL room temperature Accutase (Gibco) to detach the cells from the plate. After incubation, the ECs were collected and magnetically enriched using the CD34 microbead kit and MACS columns (Miltenyi Biotec) according to the manufacturer’s protocol. The purified cells (purity determined as mentioned below) were then plated on Animal-Component-Free coated cell culture plates using Endothelial Expansion Media (StemCell Technologies) for expansion. To harvest cells for flow cytometry, each well was aspirated of media after 3 days of the first passage, and incubated with 1 ml of room temperature Accutase for 5 minutes at 37°C to detach the cells from the plate. After incubation, 1 mL of MACs Separation Buffer (Miltenyi) was added to each well as a staining buffer to collect and transfer the detached cells to a microcentrifuge tube. The cells were then centrifuged at 300 x g for 5 minutes. The supernatant was aspirated, and each tube was stained with 5 μL of APC-Cy7 Mouse Anti-Human CD31 (PECAM-1) antibody (clone WM59, BD Biosciences) and 5 μL of APC anti-human CD144 (VE-Cadherin) antibody (clone BV9, BioLegend) to assess EC purity. The cells were then incubated in the dark for 20 minutes at 4°C. After the incubation, the cells were centrifuged at 300 x g for 5 minutes and the supernatant was aspirated. Flow cytometry was performed on a Cytoflex Flow Cytometer (Beckman Coulter) and analyzed using FlowJo Version 10.9.0 software.

#### CM Differentiation

PSC-derived CMs were differentiated from all the PSC lines implementing the Alternate GiWi protocol, using small molecule temporal modulation of the Wnt pathway, as previously described in Lian et al.^40^ Prior to differentiation, PSCs were maintained in TeSR E8 media on Matrigel-coated cell culture plates until they reached full confluency. Following 17 days of differentiation, CMs cultured on Matrigel-coated cell culture plates were harvested and stained for Anti-Cardiac Troponin T (clone 1C11) Antibody (Abcam) and Goat anti-Mouse IgG (H+L) Cross-Adsorbed Secondary Antibody Alexa Fluor 647 (Thermo Fisher Scientific) to assess CM purity as described by Zhang et al.^39^ Flow cytometry was performed on a Cytoflex Flow Cytometer (Beckman Coulter) and analyzed using FlowJo Version 10.9.0 software.

#### cFib Differentiation

PSC-derived cFibs were differentiated from all the PSC lines as described in Zhang et al.^39^ Prior to differentiation, PSCs were maintained in TeSR E8 media on Matrigel-coated cell culture plates until they reached full confluency. Following 20 days of differentiation, cFibs cultured on Matrigel-coated cell culture plates were harvested and stained for Anti-Fibroblast Antibody Clone TE-7 (Millipore Sigma) and Antibody to Sarcomeric Myosin MF20 (University of Iowa Developmental Studies Hybridoma Bank) to assess cFib purity as described by Zhang et al.^39^ Flow cytometry was performed on a Cytoflex Flow Cytometer (Beckman Coulter) and analyzed using FlowJo Version 10.9.0 software.

### Fluorescent Labeling of Effector Immune Cells

Staining of the effector cells with Calcein-AM (Invitrogen) was performed in accordance to manufacturers recommendations. After counting the effector cells, the requisite number of cells required for the experiment was first aliquoted and washed with RPMI-1640 without phenol red (Gibco). After the wash, the cells were resuspended to 2-4×10^6^ cells/mL with warm (37°C) RPMI-1640 without phenol red. The cells were added with 2.98 μL of 3 μM Calcein-AM stock solution (Invitrogen) per mL of media, mixed well with a serological pipette, and incubated in the dark at 37°C for 30 minutes. Following incubation, the cells were centrifuged twice at 450 x g for 5 minutes to remove excess Calcein-AM and resuspended at 1×10^6^ cells/mL with warm RPMI-1640 without phenol red. Violet Proliferation Dye 450 (VPD450; BD Biosciences) was used as an alternative to Calcein-AM for some experiments where the cell harbored a green fluorescent protein. Effector cells were counted and the requisite number of cells for the experiment were washed twice using 1 X PBS−/− to remove any residual serum proteins. The washed cells were then resuspended at 10e6/ml of PBS−/−. 1ul of 1uM VPD450 stock was added to the cells per mL of PBS-/– and incubated for 15 minutes in the dark at 37°C. After incubation, the cells were washed (centrifuge at 300g for 5 minutes), and the cell pellet was resuspended in warm RPMI-1640 without phenol red with 10% fetal bovine serum at 1×10^6^ cells/mL.

### Leukocyte Adhesion Assays

Adhesion assays of fluorescently-labeled immune effector cells with PSC-derived cells were performed using a combination of methods described by Zhang et al.^75^ and Wilhelmsen et al.^76^ All the PSC-derived cell types (ECs, CMs, and cFibs) were grown on a 24 well plate (Corning) and stimulated with IFNγ (50ng/ml) and TNF⍺ (10ng/ml) 48 hours prior to the experiment. After incubation, cell culture supernatants were aspirated and the wells were washed twice with RPMI-1640 with 10% bovine serum albumin (wash media) at room temperature. For experiments that incorporated ICAM-1 blocking antibody, following the wash, 500ul of fresh cell culture media for each respective cell type were combined with 10 µl of 0.5ug/µl purified anti-human ICAM-1 blocking antibody (Clone HA58, BioLegend), added to the wells, and incubated at 37°C for 1 hour. Following incubation, the wells were washed once with room temperature wash media to remove excess unbound blocking antibody, followed by addition of U937s, PBMCs, or neutrophils (1e6/ml), and then incubated for 20-30 minutes. After incubation with immune cells, the wells were washed twice with cold wash media to remove excess unbound immune cells. The wells were imaged at baseline (without immune cells) and both before and after wash, with both phase contrast and fluorescent settings using Echo Revolve | R4 Microscope enabled with Echo PRO imaging software by Bioconvergence (BICO) to document the confluency and the auto-fluorescence, respectively, prior to the addition of the effector cells. A minimum of five randomly selected regions of interest were imaged for both phase contrast and fluorescence settings; the images were then analyzed with ImageJ software (version 1.54g) to determine the mean adhered effector cells to the PSC-derived cell monolayer. The exposure settings were maintained across all images.

### Luminex for Soluble ICAM-1

Luminex assays detecting soluble ICAM-1 (sICAM1) were conducted using the MILLIPLEX Human Sepsis Magnetic Bead Panel 1 Kit (Millipore Sigma) according to the manufacturer’s protocol. Assays were run on the Luminex 200 Instrument System (Millipore Sigma) utilizing xPONENT running software. Results were analyzed using Belysa 1.1.0 analysis software.

### Teratoma Assay

Two confluent wells of PSCs were harvested from a Matrigel-coated (Corning) 6-well cell culture plate. 1-2×10^6^ cells were combined with 50 μL Matrigel (Corning) and injected subcutaneously into the hind leg of female NSG-W mice (6-8 weeks old). Teratomas were permitted to form for 6-10 weeks after the initial cell injections. Animals were then sacrificed and the teratomas were collected for histology. Teratoma tissue was submitted to the UW Department of Pathology and Laboratory Medicine’s Translational Research Initiative in Pathology (TRIP) Core for histological analysis and processing for hematoxylin and eosin staining. All animal work was conducted according to relevant national and international guidelines under the approval of the UW Institutional Animal Care and Use Committee.

### HLA Typing

HLA typing was performed on both H9 and PED05 PSC lines and for PBMC3 and PBMC5 donors. The LinkSeq HLA-ABCDRDQDP+ 384 typing kit was used as previously described in Del Rio et al.^49^

### U937 Cell Line

The U937 cell line was obtained from ATCC. It exhibits monocyte morphology and was originally derived from malignant cells obtained from the pleural effusion from a patient with histiocytic lymphoma.^77^ LFA-1 (CD11a/CD18) was assessed with a FITC anti-human antibody (clone m24, Biolegend), and MAC-1 (CD11b/CD18) with a Brilliant Violet 510 antibody (clone ICRF44, Biolegend), by flow cytometry.

### PBMC and Neutrophil Isolation

PBMCs from donors “PBMC3” and “PBMC5” were obtained from leukophoresis (BioIVT). Leukopaks were processed with Lymphocyte Separation Medium (Cellgro). PBMCs were collected via centrifugation using SepMate™ PBMC Isolation Tubes (StemCell Technologies) per the manufacturer’s recommendations. Neutrophils from an unidentified donor were obtained from whole blood using EasySep™ Direct Human Neutrophil Isolation Kit (StemCell Technologies) following the manufacturer’s instructions. The Easy 50 EasySep™ Magnet (StemCell Technologies) was used to isolate the neutrophil population (verified by flow cytometry using an APC-cy7 antibody specific for human CD16 (clone 3G8, BD Biosciences) and a PE antibody for human CD66b (clone C10F5, BD Biosciences). LFA-1 and MAC-1 were assessed as described in “U937 Cell Line,” above.

### RNA Sequencing Analysis

Library preparation, sequencing, quality control, alignment, and expression estimation were performed at the UW-Madison Biotechnology Center’s Bioinformatics Core Facility (Research Resource Identifier – RRID:SCR_017769; Madison, WI, United States). Specifically, Illumina TruSeq RNA libraries were constructed and 150 bp paired-end sequencing was performed using the Illumina Hi-Seq 2500 platform. Read quality was evaluated using FastQC.^78^ Bioinformatic analysis of RNASeq reads adhered to ENCODE guidelines and best practices for RNASeq.^79^ Briefly, alignment of adapter-trimmed (Skewer v0.1.123)^80^ 2 × 150 bp paired-end strand-specific Illumina reads to the GRCh38.p14 genome was achieved with the Spliced Transcripts Alignment to a Reference (STAR v2.5.3a) software,^81^ and a splice-junction aware aligner using Ensembl annotation.^82^ Expression estimation was conducted using RSEM v1.3.0 (RNASeq by Expectation Maximization).^83^ To test for differential expression among individual group contrasts, expected read counts were used as input into edgeR v3.16.5.^84^ Significance of the negative-binomial test was adjusted with a Benjamini–Hochberg false discovery rate correction at the 5% level.^85^ Before statistical analysis with edgeR, independent filtering was performed, requiring a threshold of at least 1 read per million in two or more samples.

### Mixed Lymphocyte Reaction

Mixed lymphocyte reactions were performed similar to our previously published method.^86^ Briefly, highly pure (>85% cTNT^+^) PSC-CMs were made from H9 WT and ICAM-1 KO PSCs, as described above. Cells were co-cultured for 6 days with HLA-mismatched PBMCs (1:6 Target:Effector ratio) that had been labeled with VPD450 proliferation dye, as described above. Alloreactivity was assessed via proliferation (VPD450 dye dilution) of total viable cells and various T cell subpopulations by flow cytometry. Total CD8^+^CD3^+^ T cell and effector memory T (TEM) cell (CD45RO ^+^ CD45RA^-^CD62L^-^CCR7^-^) proliferation was determined by staining with the following antibodies (clones UCHL1, HI100, DREG-56, G043H7, respectively), as well as excluding dead cells with LIVE/DEAD Fixable Red stain (Thermo Fisher Scientific), to determine the percentage of cells that had diluted VPD450 and thus proliferated. Relative proliferation index was calculated by using the ratio of the percentage of proliferating cells with PSC-CM targets divided by the baseline proliferating cells without targets. N=3 biological replicates, error bars = standard deviation, analysis via 2-tailed unpaired t-test, Prism 9.3.1.

### BLI in NeoThy Humanized Mice

NOD.Cg-KitW-41J Tyr + Prkdcscid Il2rgtm1Wjl/ThomJ (NBSGW) mice^50^ (6 to 8 weeks old) were obtained from the Jackson Laboratory. NeoThy humanized mice were made, as previously described,^49^ with neonatal cord blood and thymus tissue. Following verification of human immune cell chimerism (>10% human CD45^+^ and >10% human CD3^+^, by flow cytometry, as described above, using clones HI30 and UCHT1, respectively), the WT and ICAM-1 KO hPSC lines (PED05ALG at passage 47 and PED05 CD54KO ALG at passage 54, respectively) cultured as described above, were injected intramuscularly into the hind limb of the animals with Matrigel. Akaluc reporter signal was assessed via BLI using the IVIS Spectrum Imaging System (PerkinElmer) at three timepoints following the injections. AkaLumine substrate solutions were prepared dissolving 10mg AkaLumine-HCl (AOBIOUS) in 1 mL of PBS−/− (Corning), and mice were injected intraperitoneally with 100 μL. Animals were anesthetized using isoflurane and imaged 30 minutes after injection. To account for variations in peak signal intensity time, images were acquired sequentially once every minute for 5 minutes, and the images with the highest signal were selected for analysis. Images were analyzed using Living Image 4.7 software (PerkinElmer), and maximum regions of interest were quantified in units of photons per second.

### Statistical Analysis and Graphing

GraphPad Prism (version 10.2.2) was used for statistical analysis and plotting of graphs for all experiments, except RNA sequencing, as indicated above. For leukocyte binding assays, one-way ANOVA assays with Tukey’s multiple comparisons test were used to determine statistical significance.

## Acknowledgments

This study was supported in part by the Wisconsin Alumni Research Foundation (WARF), NIH NHLBI R56HL165189 (M.E.B.), NIH NIAID 75N93021C00004 (M.E.B), NIH NHLBI U01HL134764 (T.J.K., M.E.B.), the UW Carbone Cancer Center Support Grant P30 CA014520, the UW Stem Cell and Regenerative Medicine Center Graduate Research Fellowship (S.S.), and NIH NIDOCD T32DC009401 (J.S.). The authors thank Jennifer Zellner for expert manuscript editing assistance; Andres Mejia for helpful discussions regarding teratoma pathology; Finn McCarthy for assistance with literature review; the UW Biotechnology Gene Expression Center (Research Resource Identifier – RRID:SCR_017757) for library preparation and the DNA Sequencing Facility (RRID:SCR_017759) for sequencing; Anita Bhattacharyya and Andrew Petersen at the UW Human Stem Cell Core for assistance with CRISPR/Cas9 guide RNA design and editing (supported in part by a core grant to the Waisman Center from NIH NICHD P50HD105353); Jenny Gumperz for helpful discussion and Dana Baiu for neutrophil isolation assistance; Jinhua Zhang for advice related to cardiac fibroblast differentiation; Seth Taapken and WiCell for karyotyping and quality-control reagents; and the UW Translational Research Initiative in Pathology (TRIP) Histology lab. Graphical Abstract and Figure 2A were created with BioRender.com.

