## Supplementary Figures and Table for "Diminished Immune Cell Adhesion in Hypoimmune ICAM-1 Knockout Pluripotent Stem Cells"

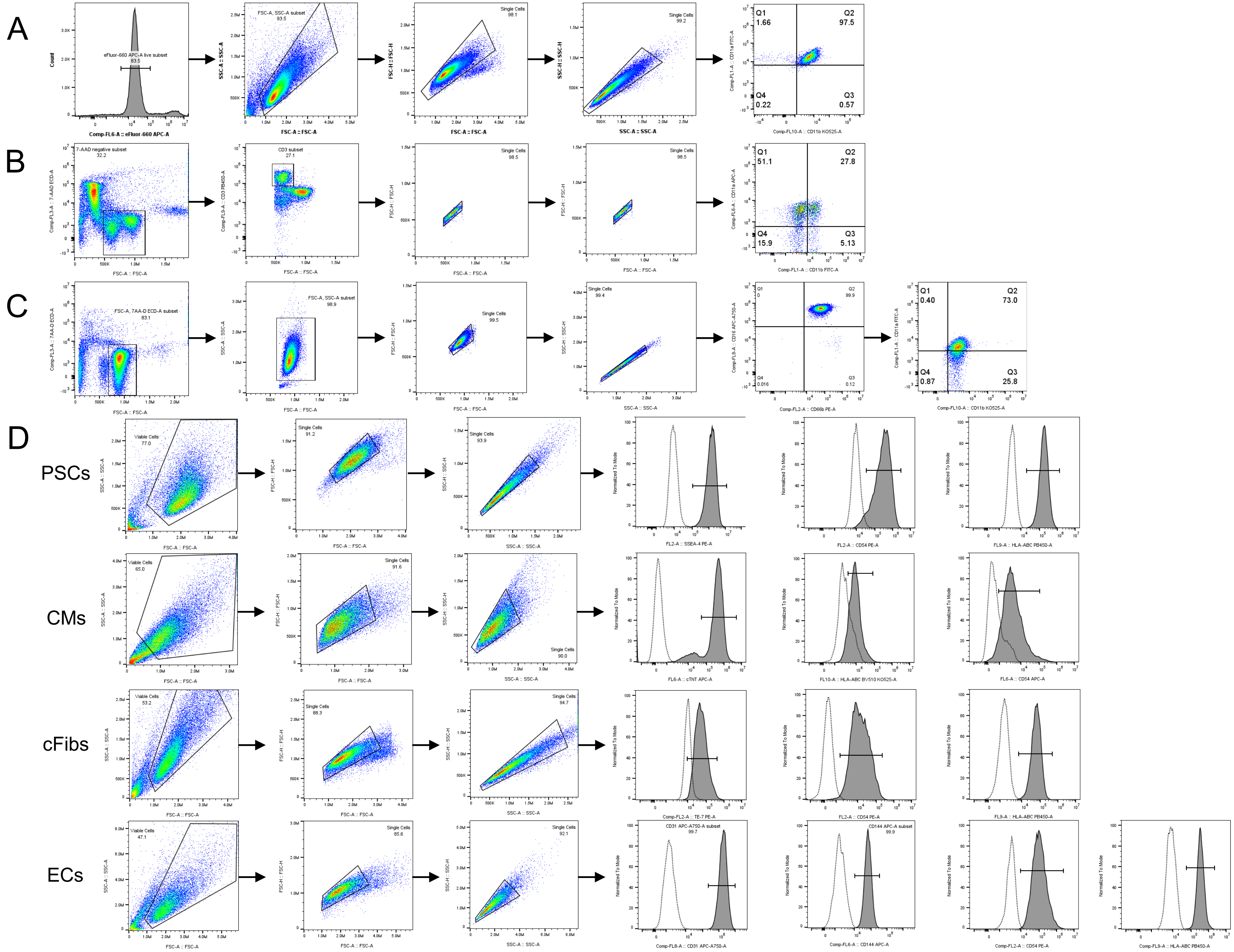


**Supplementary Figure 1. Flow Cytometry Gating Strategies.** CD18 forms complexes with CD11a (i.e., LFA-1) and CD11b (i.e., MAC-1), which are present at differing levels on all innate and adaptive immune cells within peripheral blood mononuclear cells (PBMCs). We stained effector cells including (A) U937 monocytic cell line, (B) PBMCs, and (C) neutrophils with anti-CD11a and CD11b antibodies. All the effector cells were stained with various viability stains, as indicated below, to determine the live cell population gate. (A) U937s were stained with eFluor660 Live/Dead viability dye to select the eFluor-negative live cell fraction to assess the surface CD11a and CD11b. (B) PBMCs were stained with 7-AAD. The 7-AAD-negative fraction (i.e., live PBMCs) was first gated upon, followed by gating on the CD3-positive fraction. After the removal of the doublets from the CD3-positive fraction, the live population of singlets was then assessed for surface CD11a and CD11b. (C) The isolated primary human neutrophils were stained with 7-AAD and the negative fraction was gated upon to select the live cells. From the live cells, the granulocyte fraction (nearly 99%) was then gated upon followed by the removal of doublets. The singlets were then assessed for CD16 and CD66b (surface markers of neutrophils) and the double positive population (nearly 100%) was then assessed for surface CD11a and CD11b. The gates are drawn based on the unstained control (data not shown). (D) Gating strategy for pluripotent stem cells (PSCs) and PSC-derived cell types: No live/dead stain was used for PSC and PSC-derived cells because of typical good survival rate from cell cultures (data not shown). All PSCs and PSC-derived cell types were first removed of doublets. The singlet population was then assessed for surface CD54 and HLA-ABC. The singlet population was also assessed for distinct surface markers such as pluripotency marker Stage-specific Embryonic Antigen-4 (SSEA-4) for PSCs, Cardiac Troponin T (cTnT) to assess the purity of cardiomyocytes (CMs), Fibroblast TE-7 to assess the purity of cardiac fibroblasts (cFibs), in addition to CD31 and CD144 to assess the purity of endothelial cells (ECs). The dotted lines represent the unstained (negative) control population and the dark grey peaks represent the stained population in the histograms. Analysis was performed via FlowJo 10.10 software. Biological replicates = 3.

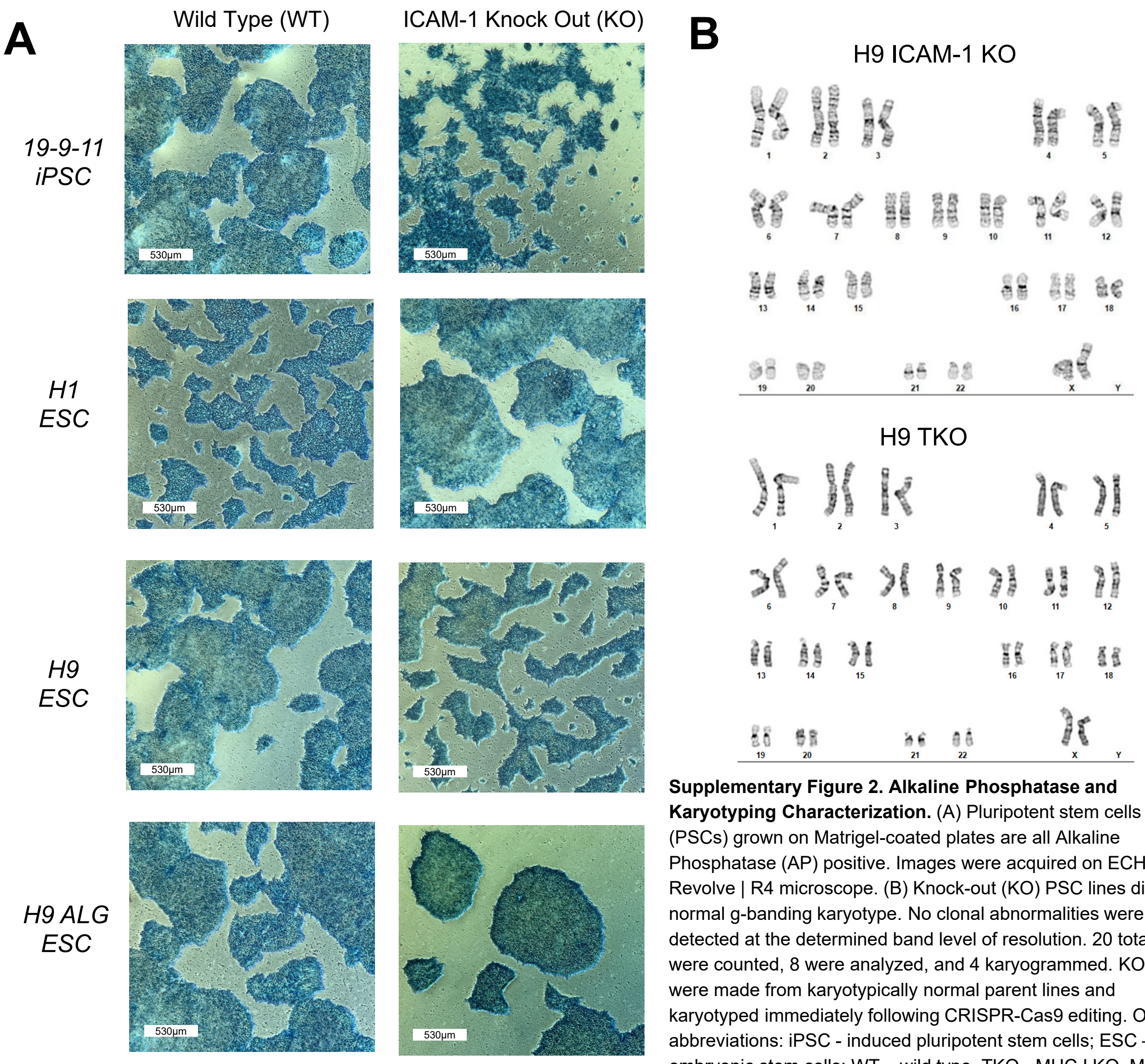


**Supplementary Figure 2. Alkaline Phosphatase and Karyotyping Characterization.** (A) Pluripotent stem cells (PSCs) grown on Matrigel-coated plates are all Alkaline Phosphatase (AP) positive. Images were acquired on ECHO Revolve | R4 microscope. (B) Knock-out (KO) PSC lines display a normal g-banding karyotype. No clonal abnormalities were detected at the determined band level of resolution. 20 total cells were counted, 8 were analyzed, and 4 karyogrammed. KO lines were made from karyotypically normal parent lines and karyotyped immediately following CRISPR-Cas9 editing. Other abbreviations: iPSC - induced pluripotent stem cells; ESC – embryonic stem cells; WT – wild type. TKO - MHC I KO, MHC II KO, ICAM-1 KO, CD47-overexpressing hPSC line.

| | *HLA-Class I* | | | | *HLA-Class II* | |
|---|---|---|---|---|---|---|
| | **A** | | **B** | | **DRB** | |
| **PED05** | 1 | 2 | 8 | 51 | 11 | 17 |
| **PBMC3** | 30 | 30 | 7 | 7 | 9 | 9 |
| **PBMC5** | 2 | 3 | 44 | 47 | 4 | 12 |
| **H9 PSC line** | 2 | 3 | 35 | 44 | 15 | 16 |

**Supplementary Table 1. Donor Peripheral Blood Mononuclear Cells (PBMC) and Pluripotent Stem Cell (PSC) Cell Line HLA Typing.** HLA Class I and Class II typing is shown for PBMC donors and PSC lines used for the experiments included in the manuscript.

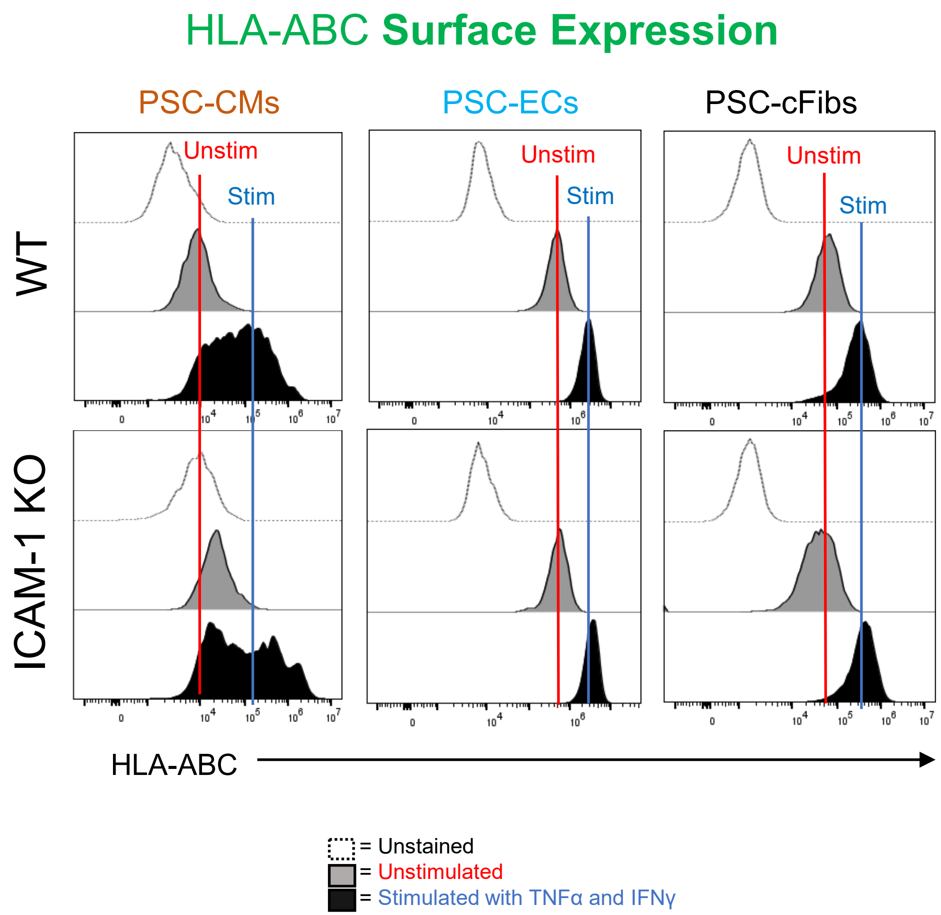


**Supplementary Figure 3. HLA-ABC Surface Expression on Multiple Differentiated Cell Types.** H9 pluripotent stem cells (PSCs) were differentiated and expanded into highly pure cardiomyocytes (CMs; left), endothelial cells (ECs; middle), and cardiac fibroblasts (cFibs; right) and stimulated with 10 ng/ml TNFα and 50 ng/ml IFNγ for 48 hours. Cells were harvested and stained with HLA-ABC antibody for flow cytometry. HLA-ABC (Class I) upregulation was assessed in both wild type (WT; top) and ICAM-1 knock-out (KO; bottom) cells. Analysis was performed via FlowJo 10.10 software. Data are representative of n=3 experiments.

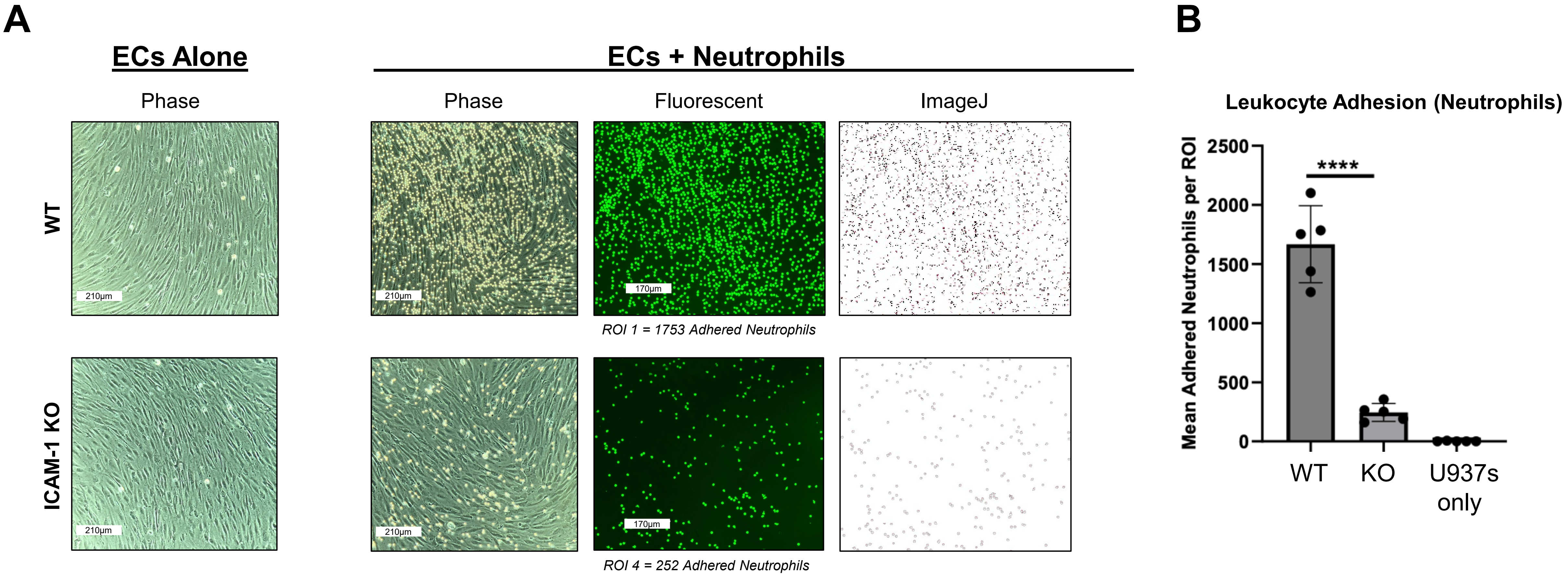


**Supplementary Figure 4. Neutrophil Adhesion Assay with ICAM-1 Knock-out (KO) Pluripotent Stem Cell (PSC)-Derived Endothelial Cells (ECs).** H9 wild type (WT) and ICAM-1 KO PSCs were differentiated into high purity (CD31+CD144+) ECs and stimulated with TNFα and IFNγ for 48 hours, similar to Figure 6. Primary human neutrophils were cultured with the WT and ICAM-1 KO ECs and washed. Images of neutrophil adhesion to the stimulated ICAM-1 KO ECs were compared to the stimulated WT control ECs for analysis. Images were acquired on an ECHO Revolve | R4 microscope (10X), and analyzed using ImageJ version 1.54g. **** = $p<0.0001$, biological replicates = 3. Statistical analysis was performed via Prism 10.2.2 software. ROI = Region of Interest

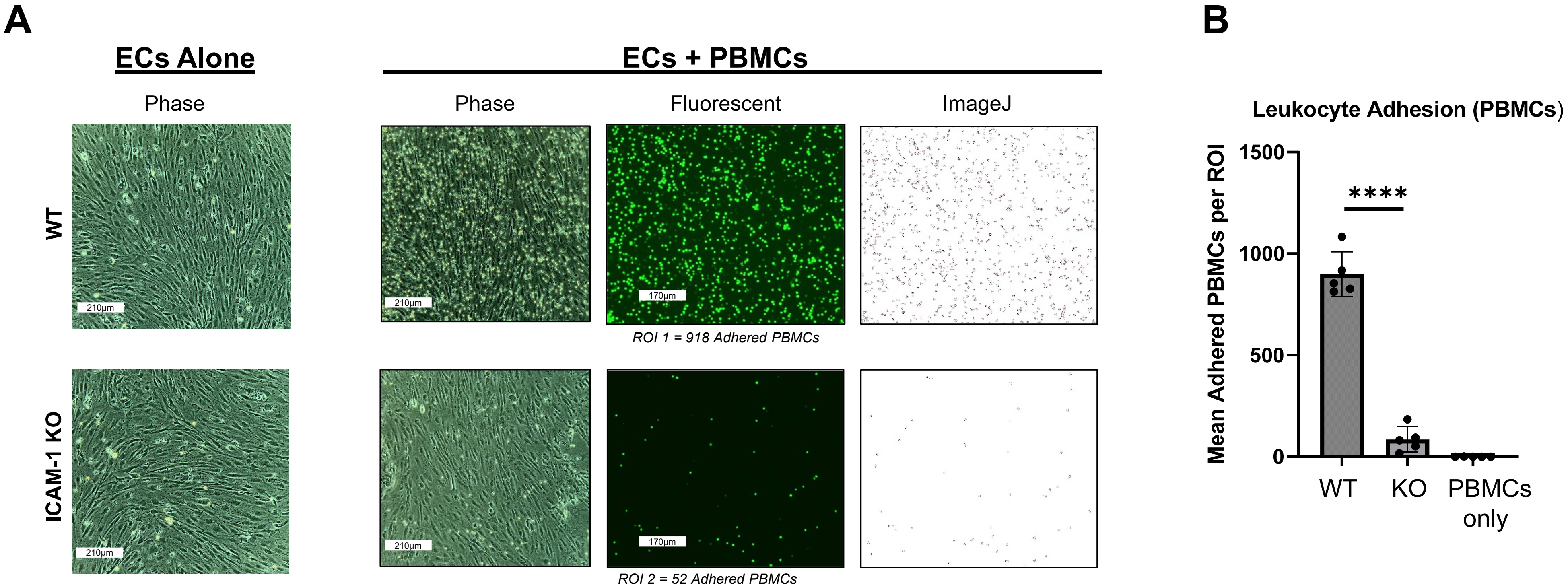


**Supplementary Figure 5. Peripheral Blood Mononuclear Cell (PBMC) Adhesion Assay with ICAM-1 Knock-out (KO) Pluripotent Stem Cell (PSC)-Derived Endothelial Cells (ECs).** H9 wild type (WT) and ICAM-1 KO PSCs were differentiated into high purity (CD31+CD144+) ECs and stimulated with TNFα and IFNγ for 48 hours, similar to Figure 6 and Supplementary Figure 4. PBMCs were cultured with the WT and ICAM-1 KO ECs and washed. Images of PBMC adhesion to the stimulated ICAM-1 KO ECs were compared to the stimulated WT control ECs for analysis. Images were acquired on an ECHO Revolve | R4 microscope (10X), and analyzed using ImageJ version 1.54g. **** = $p<0.0001$, biological replicates = 3. Statistical analysis was performed via Prism 10.2.2 software. ROI = Region of Interest

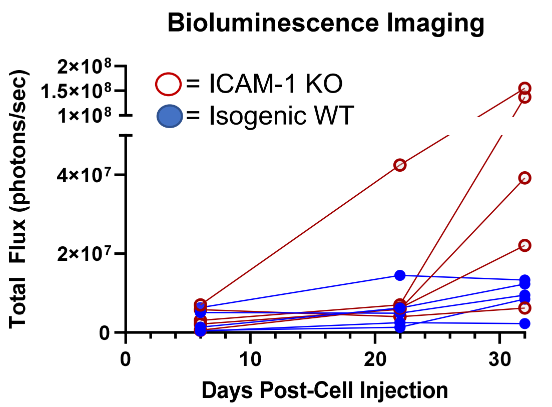


**Supplementary Figure 6. Transplantation of ICAM-1 Knock-out (KO) and Wild Type (WT) Pluripotent Stem Cells (PSCs) into Humanized Mice.** ICAM-1 KO PSCs (1x106 cells) harboring a constitutively expressed Akaluc reporter were co-injected with Matrigel into the right leg of NeoThy humanized mice (>10% hCD3+) with HLA-mismatched immune systems. Isogenic WT PSCs harboring a constitutively expressed Akaluc reporter were co-injected with Matrigel into the left leg of the same mice. Bioluminescence imaging of engraftment-positive cells was conducted at multiple time points for 32 days to monitor retention of the graft in the in vivo presence of human immune effector cells, as indicated by the persistence of total flux signal.

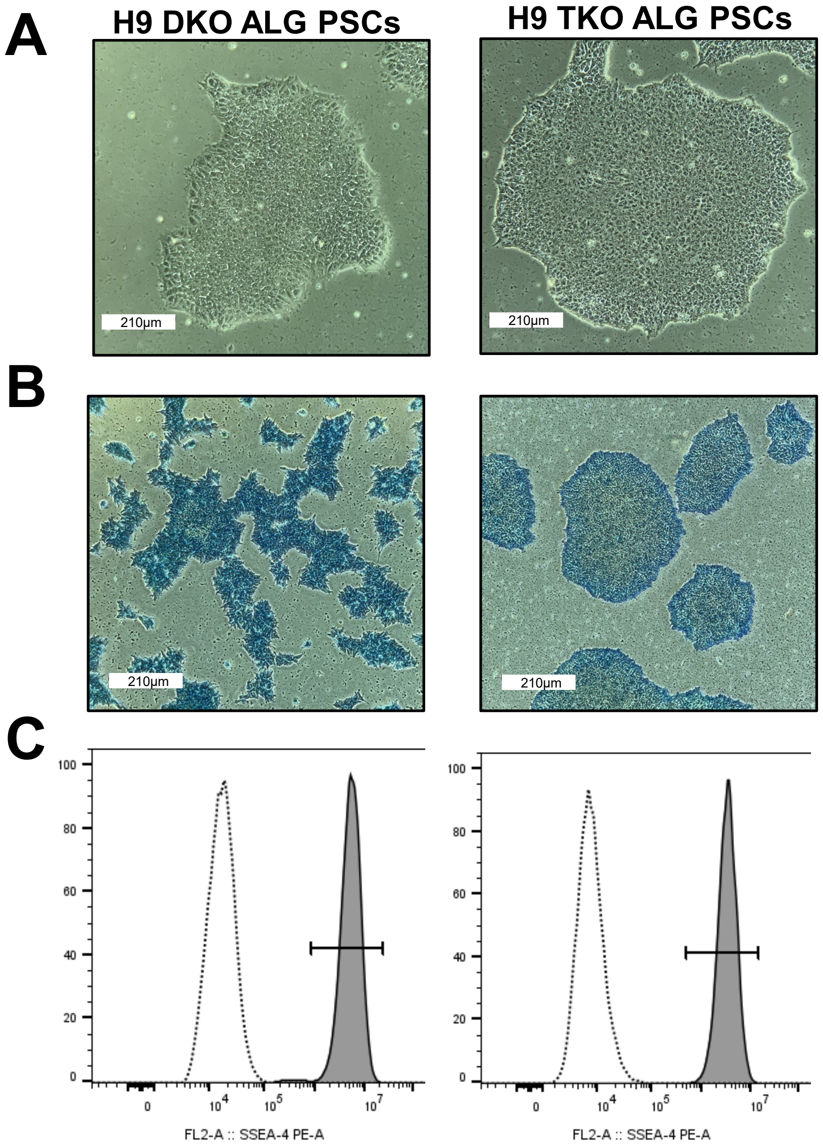


**Supplementary Figure 7. Morphology and Pluripotency Characterization of Triple Knock-out (TKO) Hypoimmune Pluripotent Stem Cell (PSC) Lines.** (A) Phase contrast images (10X) of "first generation" H9 PSC line including knock-out (KO) of β2M and CIITA plus overexpression of CD47, also harboring constitutive Akaluc + GFP (ALG) construct ("DKO"). An ICAM-1 KO was added to the DKO PSC line, creating the "TKO" PSC line. (B) Phase contrast images were acquired (4X) after alkaline phosphatase. (C) SSEA-4 pluripotency staining on both of the cell lines. Data are representative of n=3 experiments. Images were acquired on ECHO Revolve | R4 microscope, flow cytometric data analysis was performed via FlowJo 10.10 software.
